# An IFN-STAT1-CYBB Axis Defines Protective Plasmacytoid DC to Neutrophil Crosstalk During *Aspergillus fumigatus* Infection

**DOI:** 10.1101/2024.10.24.620079

**Authors:** Yahui Guo, Mariano A. Aufiero, Kathleen A.M. Mills, Simon A. Grassmann, Hyunu Kim, Paul Zumbo, Mergim Gjonbalaj, Audrey Billips, Katrina B. Mar, Yao Yu, Doron Betel, Joseph C. Sun, Tobias M. Hohl

## Abstract

*Aspergillus fumigatus* is the most common cause of invasive aspergillosis (IA), a devastating infection in immunocompromised patients. Plasmacytoid dendritic cells (pDCs) regulate host defense against IA by enhancing neutrophil antifungal properties in the lung. Here, we define the pDC activation trajectory during *A. fumigatus* infection and the molecular events that underlie the protective pDC - neutrophil crosstalk. Fungus-induced pDC activation begins after bone marrow egress and results in pDC-dependent regulation of lung type I and type III IFN levels. These pDC-derived products act on type I and type III IFN receptor-expressing neutrophils and control neutrophil fungicidal activity and reactive oxygen species production via STAT1 signaling in a cell-intrinsic manner. Mechanistically, neutrophil STAT1 signaling regulates the transcription and expression of *Cybb*, which encodes one of five NADPH oxidase subunits. Thus, pDCs regulate neutrophil-dependent immunity against inhaled molds by controlling the local expression of a subunit required for NADPH oxidase assembly and activity in the lung.

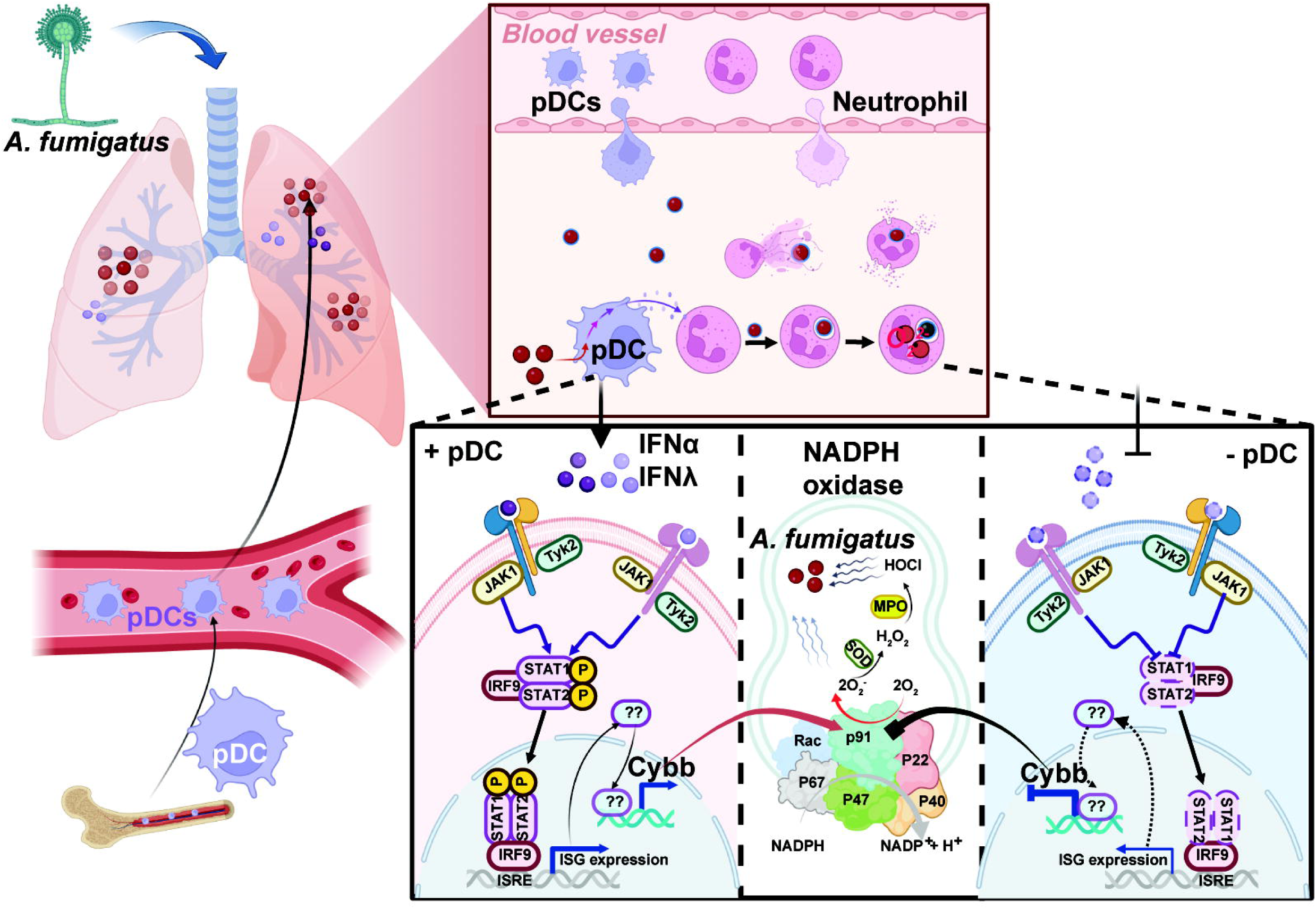

## Introduction

Invasive pulmonary aspergillosis, a life-threatening mold infection, occurs when the respiratory immune system fails to eradicate ubiquitous inhaled *Aspergillus* spores (i.e., conidia) prior to their germination into tissue-invasive hyphae (1). At-risk populations include patients with acute leukemia and other bone marrow disorders, recipients of hematopoietic cell and lung transplants, individuals with immune-related or neoplastic diseases treated with prolonged corticosteroid therapy or with novel targeted biologics (e.g., ibrutinib) that blunt fungal immune surveillance pathways (2–4) and patients with severe respiratory virus infections, including influenza and COVID-19 (5, 6). As a result of these medical advances and global pandemic viruses, *A. fumigatus* has become the most common agent of mold pneumonia worldwide (6–12).

Host defense against airborne mold conidia depends on intact myeloid cell numbers and function at the respiratory mucosa. Lung-infiltrating neutrophils and monocyte-derived dendritic cells (Mo-DCs) play essential roles in killing phagocytosed conidia, with a central role for products of NADPH oxidase in this process (3, 13–15). Patients with chronic granulomatous disease (CGD) and defective NADPH oxidase function are uniquely vulnerable to invasive aspergillosis (IA, with a lifetime prevalence of 40-55% (16). Exposure to products of NADPH oxidase induces a regulated cell death process in conidia trapped within neutrophil phagosomes, resulting in sterilizing immunity at respiratory mucosal barrier (17).

An emerging theme of host defense against *Aspergillus* is the essential role of intercellular crosstalk to license neutrophil effector properties *in situ*. On one hand, recruited monocytes and Mo-DCs promote neutrophil ROS production through type I and III interferon (IFN) release, though the link between IFN production and neutrophil ROS activity remains undefined (15). On the other hand, we discovered that fungus-engaged neutrophils and Mo-DCs release CXCL9 and CXCL10 which results in the recruitment of CXCR3^+^ pDCs by promoting their influx from the circulation into the lung. In the lung, pDCs enhance neutrophil fungicidal properties and are essential for host defense, even in the presence of lung-infiltrating monocytes and Mo-DCs (18). However, the underlying molecular mechanisms that regulate pDC to neutrophil crosstalk remain undefined. Thus, both Mo-DCs and pDCs independently enhance neutrophil antifungal activity, but it is unknown whether pDCs employ either similar or distinct mechanisms of intercellular crosstalk with neutrophils compared to monocytes and Mo-DCs. Critical open questions relate to the identity of pDC-derived molecules that are required for protective crosstalk with neutrophils and to the ensuing molecular changes in neutrophils that regulate antifungal activity *in situ*.

In this study, we demonstrate that pDCs represent a key and indispensable source of type I and type III interferons (IFNs) in the lung during respiratory *A. fumigatus* infection. In turn, pDC-dependent and IFN-induced STAT1 signaling controls neutrophil *Cybb* expression, which encodes an essential subunit of the NADPH oxidase complex. Thus, protective pDC to neutrophil crosstalk primarily harnesses intercellular IFN-STAT1 signaling to calibrate the synthesis of a critical component of the NADPH oxidase complex, thereby licensing neutrophils to achieve optimal antifungal activity and promoting sterilizing responses against inhaled mold spores.

## Results

### *Aspergillus fumigatus* induces pDC activation in the lung

In response to respiratory fungal infection, pDCs exit the bone marrow (BM), enter the circulation, and traffic to the lung. To gain an understanding of the pDC activation trajectory during this infection-induced trafficking event, we conducted an unbiased transcriptome analysis of BM pDCs isolated from uninfected mice and of BM and lung pDCs isolated from mice at 72 h post-infection (pi), since this time point represents the peak of lung pDC influx (Fig. 1A, 1B, Supplemental Fig. S1A). For each pDC RNA-seq sample, we pooled sorted pDCs from 10 mice from each tissue examined and analyzed 4 biological replicates. The bulk pDC transcriptome was remarkably similar between naïve and infected BM pDCs, with only 51 differentially expressed genes, supporting a model in which pDC activation occurs en route to the site of infection. In these two groups of BM pDCs, we did not observe notable differences in the transcription of genes that encode cytokines or interferons (Fig. 1C, Supplemental Fig. S1B). In contrast, RNA-seq analysis revealed significant changes in the pDC transcriptome when lung pDCs were isolated from *Aspergillus*-infected mice and compared to BM pDCs from the same animals (Fig. 1B). Overall, pDCs from infected lungs had 4475 and 4474 differentially regulated genes compared to pDCs isolated from the BM of naïve and infected mice, respectively. (Fig.1C, Supplemental Fig. S1C and S1D). Using Kyoto Encyclopedia of Genes and Genomes (KEGG) pathway enrichment analysis, we found that pathways involved in the cytokine-cytokine receptor interaction, Toll-like receptor, RIG-I, and JAK-STAT signaling were among the most upregulated pathways in pDCs isolated from infected lungs compared to BM pDCs isolated from the same mice (Fig. 1D and Supplemental Fig. S1E).

**Figure 1.**
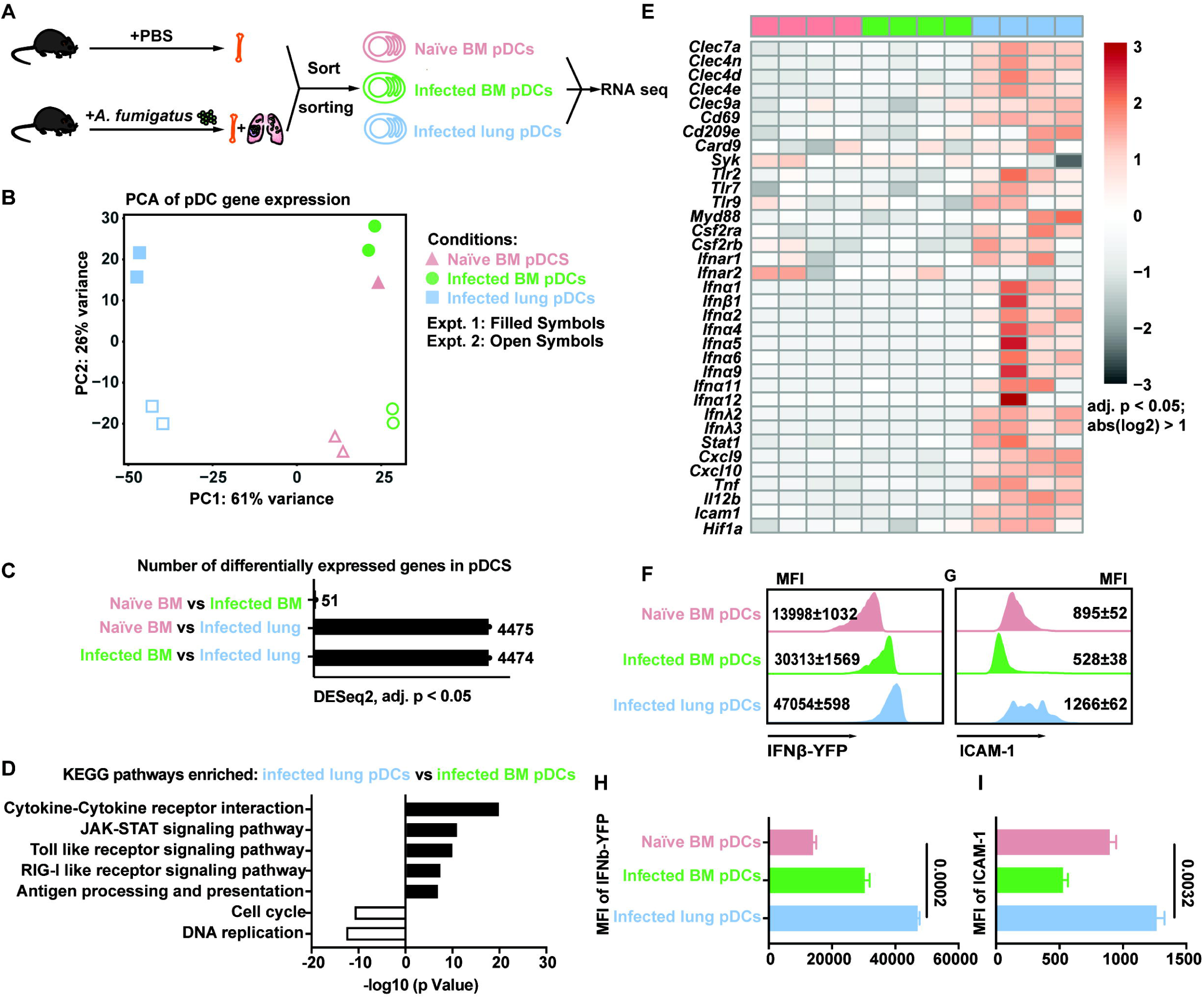
pDC transcriptome following *A. fumigatus* infection. (A) Experimental scheme for bulk RNA-seq of BM pDCs sorted from naïve mice (red symbols) and BM (green symbols) and lung (blue symbols) pDCs sorted from *A. fumigatus*-infected mice. (B) Principal component analysis of gene expression in sorted BM pDCs from naïve (triangles) and infected mice (dots) and of sorted lung pDCs from infected mice (squares). Each symbol represents a biological replicate. pDCs from 10 mice were pooled for each replicate, and 2 replicates were included in each of 2 experiments, denoted as Expt1 (filled symbol) and Expt 2 (open symbols). (C) Number of differentially expressed genes for three comparisons of 2 pDC subsets. (D) Differentially enriched KEGG pathways (q < 0.05) in pDCs isolated from infected lungs vs infected BM. Black bars indicate pathways enriched in lung pDCs from infected mice, white bars indicate pathways enriched in BM pDCs from infected mice. (E) Heatmap for 35 selected genes with a >2 -fold difference in expression and a false discovery rate (FDR) p < 0.05. Each lane represents 1 replicate from 2 expts. (F - I) Representative flow cytometry plots (F and G) and quantified mean fluorescence intensity MFI (H and I) of IFNβ YFP and ICAM1 expression in pDCs isolated from (top) naïve BM, (middle), the BM of *A. fumigatus*-infected mice or (bottom) the lungs of *A. fumigatus*-infected mice. (A-I) Infection dose: 3 × 10^7^ CEA10 conidia via intratracheal route, analysis 72 hpi. (B - E) Data were pooled from 2 independent experiments. (H and I) Statistical analysis: Kruskal-Wallis test. See also Figure S1.

Lung-infiltrating pDCs upregulated genes implicated in fungal recognition, including the C-type lectin receptors *Clec7a*, *Clec4n*, *Clec4d*, *Clec4e*, *Clec9a*, *Cd69* and *Cd209e* and downstream signaling molecules (*Card9* and *Syk*). pDCs upregulated Toll-like (*Tlr2*, *Tlr7*, *Tlr9*, and *Myd88)*, and growth factor (*Csf2ra*, *Csf2rb*) signaling pathways as well. Notably, we found that expression of type I interferon (IFN), type III IFN, and the type I IFN receptor (*Ifnar1* and *Ifnar2*), were markedly increased in lung-infiltrating pDCs isolated from infected mice (Fig. 1E), consistent with prior reports that *A. fumigatus* infection induces type I and type III IFN release in the lung (15). In addition, lung-infiltrating pDCs upregulated cytokine and chemokine (*Cxcl9*, *Cxcl10* and *Il12b)*, integrin receptor (i.e., *Icam1*, Intercellular Adhesion Molecule) and *Hif1a* mRNAs (Fig. 1E).

To examine the impact of pDC transcriptional changes on protein expression, we infected IFN*β* reporter mice with *A. fumigatus* and found that *ifnb* promoter-driven fluorescent protein expression increased when BM pDCs trafficked to the lung (Fig. 1F and 1H). Similarly, we observed pDC trafficking-dependent increases in ICAM1 surface expression (Fig. 1G and 1I). Thus, *A. fumigatus* infection substantially alters the pDC transcriptome at the portal of infection.

### pDCs regulate type I and type III IFN in the lung during *A. fumigatus* infection

To examine the contribution of pDCs to the lung inflammatory milieu, we examined type I (*Ifna1/2/5/6*) and type III (*Ifnl2/3*) IFN induction and found that induction peaked at 48 to 72 hpi (Fig. 1E and Fig. 2A-2C), temporally coincident with the peak lung pDC influx observed in a prior study (18). To determine whether pDCs directly control type I and type III induction, we infected BDCA2-DTR (pDC Depleter) mice, in which diphtheria toxin (DT) administration specifically ablates pDCs at rest and under inflammatory conditions (19), but does not ablate - CC-chemokine receptor 2 (CCR2)-expressing monocytes and Mo-DCs that have been implicated in type I and type III IFN release following *A. fumigatus* infection (15, 18). pDC depleter mice exhibited a 60-80% reduction in lung *Ifna1/2/5/6* and *Ifnl2/3* mRNA levels at 72 hpi, demonstrating that pDCs directly control the induction of type I and III IFN mRNA, consistent with the bulk RNA-seq data and with their activation trajectory in the *Aspergillus*-infected lung (Fig. 1E).

**Figure 2.**
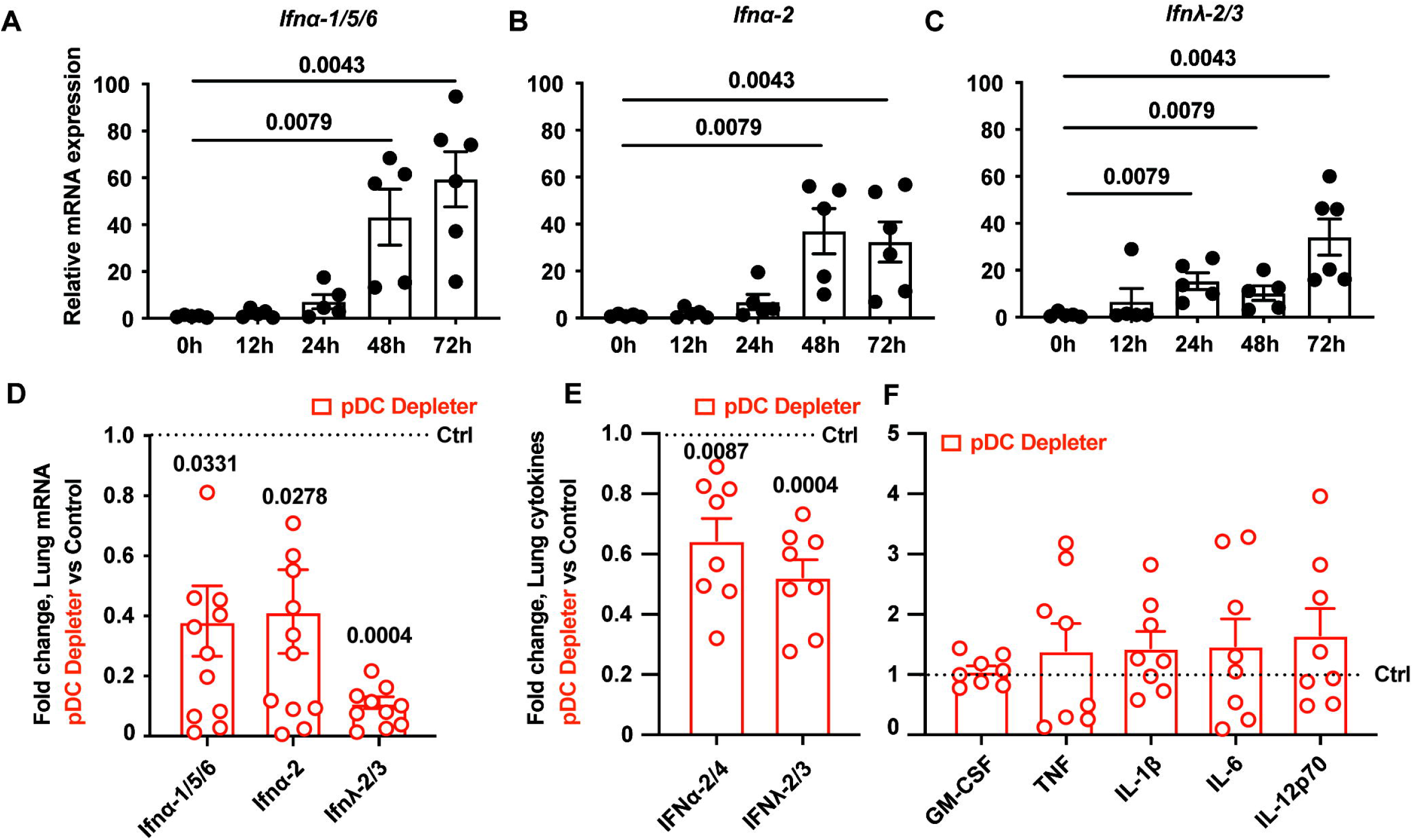
Cytokine profiles in pDC-depleted mice during *A. fumigatus* infection. (A - C) *Ifn* gene expression, measured by qRT-PCR using TaqMan probes, in the lung of C57BL6/J mice at indicated times, n = 5-6 per group. (D) Lung *Ifn* gene expression and (E and F) lung cytokine levels measured by ELISA in DT-treated pDC depleter mice (BDCA2-DTR^Tg/+^; red symbols) and DT-treated non-Tg littermate controls (black dashed line), n = 8 per group. (A - E) Infection dose: 3 × 10^7^ CEA10 conidia via intratracheal route, analysis 72 hpi. Data were pooled from 2 independent experiments and presented as mean ± SEM. Dots represent individual mice. Statistical analysis: Kruskal-Wallis test.

To measure the importance of pDCs for lung cytokine levels, we next compared lung cytokine profiles by ELISA from pDC-depleted mice and from non-transgenic, co-housed littermate controls. pDC ablation resulted in a partial depletion (40-60%) of lung type I (IFN*α* 2/4) and type III IFN levels (IFNλ 2/3). In contrast, other pro-inflammatory cytokines implicated in pulmonary antifungal defense (GMCSF, TNF, IL1β, IL6, and IL12) were not impacted by pDC ablation, due to their production by other cellular sources (12) in the lung (Fig. 2E and 2F). These data establish that pDCs play a critical role in regulating lung type I and type III IFN levels during *A. fumigatus* infection.

### STAT1 signaling in neutrophils controls the intracellular killing of *Aspergillus* conidia

Type I and type III IFNs both activate STAT1 signaling in target cells and are essential for host defense against *A. fumigatus* (15). Under baseline conditions and during *Aspergillus* infection, lung neutrophils expressed both type I IFN receptor and type III IFN receptor mRNA, as judged by RNAscope analysis (Supplemental Fig. S2A and S2B), and both signal through STAT1. Targeted ablation of *Stat1* in neutrophils renders mice susceptible to invasive aspergillosis, yet it is unknown how STAT1 signaling is coupled to myeloid cell antifungal activity and whether STAT1 signaling is required for killing in a myeloid cell-intrinsic or - extrinsic manner.

To distinguish these possibilities, we generated mixed BM chimeric mice (1:1 mixture of CD45.2^+^ *Stat1*^−/−^ and CD45.1^+^ *Stat1*^+/+^ donor BM cells → lethally irradiated CD45.1^+^ CD45.2^+^ *Stat1*^+/+^ recipients) and compared the fungicidal activity of *Stat1^-/-^* and *Stat1^+/+^* leukocytes within the same lung inflammatory context (Fig. 3A). To accomplish this, we utilized fluorescent *Aspergillus* reporter (FLARE) conidia that encode a red fluorescent protein (DsRed; viability fluorophore) and are labeled with an Alexa Fluor 633 (AF633; tracer fluorophore) (20). FLARE conidia enable us to distinguish live (DsRed^+^AF633^+^) and dead (DsRed^-^AF633^+^) conidia during leukocyte interactions with single encounter resolution (Fig. 3B).

**Figure 3.**
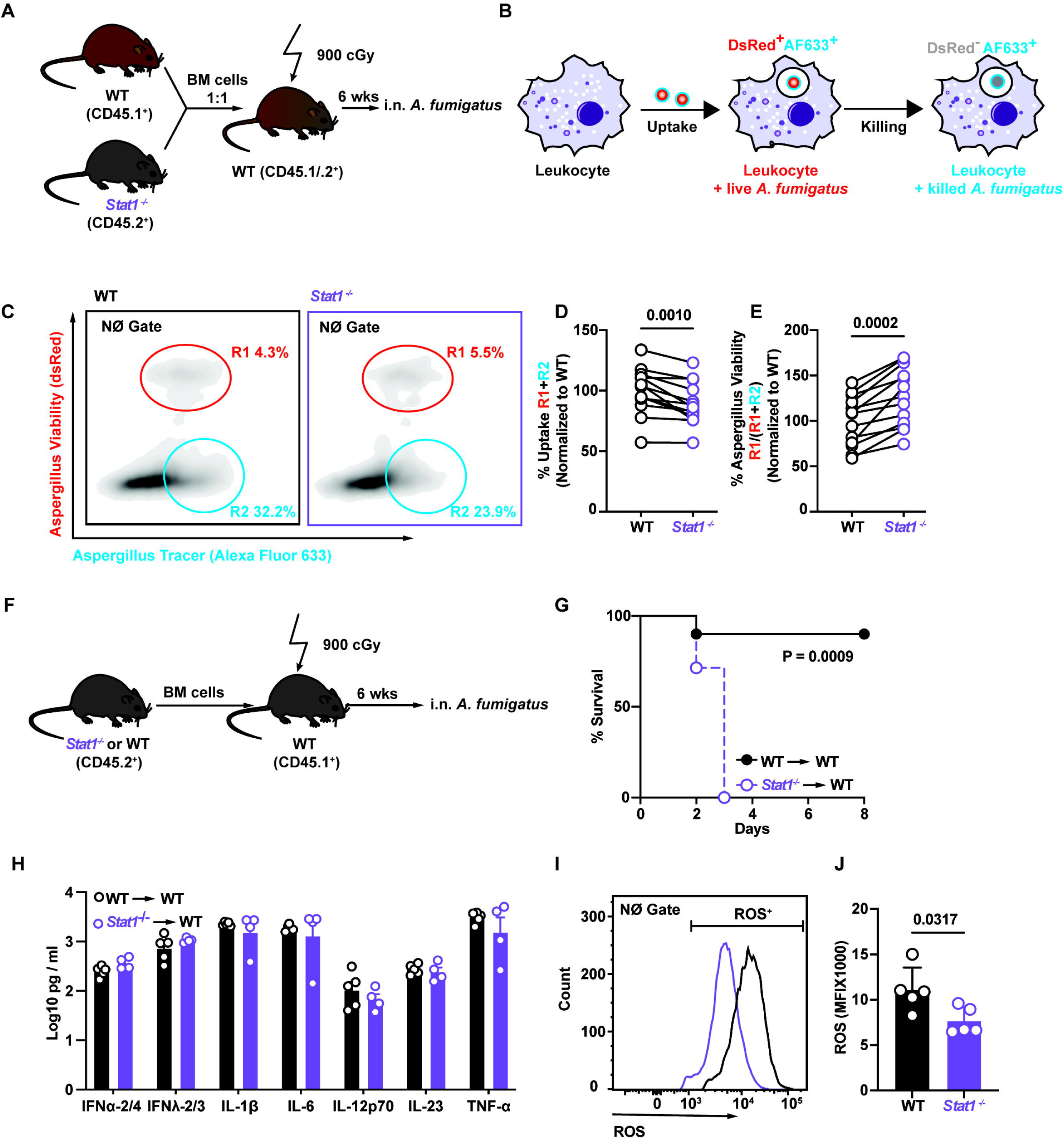
STAT1 signaling regulates neutrophil-intrinsic *Aspergillus* killing. (A) Scheme to generate *Stat1^-/-^* and *Stat^+/+^* mixed bone chimeric mice and (B) illustration of the two-component fluorescent *Aspergillus* reporter (FLARE) system used to measure conidial uptake and killing by *Stat1^-/-^*and *Stat^+/+^* lung leukocytes in mixed bone marrow chimeric mice. (C) Representative plot that display DsRed and AF633 fluorescence intensity of lung neutrophils. The R1 gate denotes neutrophils that contain live conidia, the R2 gate denotes neutrophils that contain killed conidia. (D and E) The plots show (D) normalized lung neutrophil conidial uptake (R1 + R2) ± SEM and (E) conidial viability (R1/ (R1 + R2) ± SEM in indicated lung neutrophils isolated from *Stat1^-/-^* (purple symbols) and *Stat1^+/+^*(black symbols) mixed BM chimeric mice (n=8 per group). (F) Scheme to generate *Stat1^-/-^* and WT (*Stat^+/+^*) chimeric mice. (G) Kaplan Meier survival of *Stat1^-/-^* and *Stat^+/+^*chimeric mice (n = 7-8 per group) infected with 3-6 × 10^7^ CEA10 conidia. (H) Lung cytokine levels in *Stat1^-/-^* → WT (purple symbols) and *Stat1^+/+^*→ WT (black symbols) single chimeric mice. (I) Representative plot and (J) mean ± SEM neutrophil ROS production in cells isolated from *Stat1^-/-^*→ WT (purple symbols) and *Stat1^+/+^*→ WT (black symbols) single chimeric mice (n=5 per group). (C - E) Infection dose: 3 × 10^7^ Af293 FLARE conidia via intratracheal route, analysis 72 hpi. (F - I) Infection dose: 3 × 10^7^ CEA10 conidia via intratracheal route, analysis 72 hpi. (D and E) Data were pooled from 2 independent experiments. Dots represent individual mice. Statistical analysis: paired t test. (H - J) Data are representative of 2 experiments. Dots represent individual mice. Statistical analysis: Mann-Whitney test. See also Figure S2.

Following infection with FLARE conidia, we measured the frequency of neutrophil fungal uptake (Fig. 3C, frequency of neutrophil uptake = R1 + R2) and the proportion of fungus-engaged neutrophils that contain live conidia (Fig. 3C, proportion of fungus-engaged neutrophils with live conidia = R1/ (R1 + R2)). *Stat1*^−/−^ neutrophils engulfed conidia at a slightly lower rate compared to *Stat1*^+/+^ neutrophils in the same lung (Fig. 3D). However, the frequency of fungus-engaged neutrophils that contained live conidia was higher in *Stat1*^−/−^ neutrophils compared to *Stat1*^+/+^ neutrophils (Fig. 3E). Thus, neutrophil-engulfed conidia were more likely to be viable in *Stat1*^-/-^ neutrophils than in *Stat1*^+/+^ neutrophils in the same lung tissue environment (Fig. 3E), indicating that STAT1 signaling enhances neutrophil fungicidal activity in a cell-intrinsic manner. Consistent with these findings and in line with published studies (15) we found that mice that lacked *Stat1* in radiosensitive hematopoietic cells were more susceptible to *A. fumigatus* challenge than mice with *Stat1* sufficiency in the same compartment (Fig. 3F and 3G). *Stat1* expression in radiosensitive hematopoietic cells was dispensable for lung IFN*α* 2/4, IFN*λ* 2/3, IL1β, IL6, IL12p70, IL23, and TNF levels (Supplemental Fig. S3A and 3H).

Our previous work found that lung pDCs regulate neutrophil ROS generation during respiratory *A. fumigatus* challenge (18). Here we measured neutrophil ROS production in *Stat1*^+/+^ and *Stat1*^−/−^ neutrophils and found that the ROS median fluorescence intensity (MFI) in *Stat1*^−/−^ ROS^+^ lung neutrophils were significantly reduced compared *Stat1*^+/+^ ROS^+^ lung neutrophils at 72 hpi (Fig. 3I and 3J). We enumerated myeloid cells infiltration in *Stat1*^+/+^ and *Stat1*^−/−^ lungs at 72 hpi, and found no difference in the recruitment of neutrophils, monocyte, and pDCs in *Stat1*^-/-^ mice. There was a slight decrease in Mo-DC numbers in *Stat1*^-/-^ mice, which suggests that monocytes may exhibit a limited differentiation into Mo-DCs, resulting in reduced lung Mo-DC numbers in *Stat1*^−/−^ mice compared to control mice (Fig. S3B-S3E).

### pDCs regulate neutrophil STAT1-dependent antifungal activity

To explore whether pDCs regulate neutrophil STAT1-dependent antifungal activity, we bred CD45.2^+^ BDCA2-DTR^Tg/+^ mice with CD45.2^+^ *Stat1^-/-^*mice to generate CD45.2^+^ BDCA2-DTR^Tg/+^ *Stat1^-/-^* mice and bred BDCA2-DTR^Tg/+^ *Stat1^+/+^* mice to the CD45.1^+^ (C57BL6.SJL) background. BM cells from both strains were mixed in a 1:1 ratio and utilized as donor cells to generate mixed BM chimeric mice (CD45.1^+^ BDCA2-DTR^Tg/+^ *Stat1^+/+^* and CD45.2^+^ BDCA2-DTR^Tg/+^ *Stat1^-/-^*→ CD45.1^+^CD45.2^+^ recipient mice). This experimental design enabled us to compare the fungicidal activity of *Stat1*^+/+^ and *Stat1*^-/-^ neutrophils in the same lung, either in the absence or in the presence of pDCs through the administration or omission of DT to mixed BM chimeric mice (Fig. 4A).

**Figure 4.**
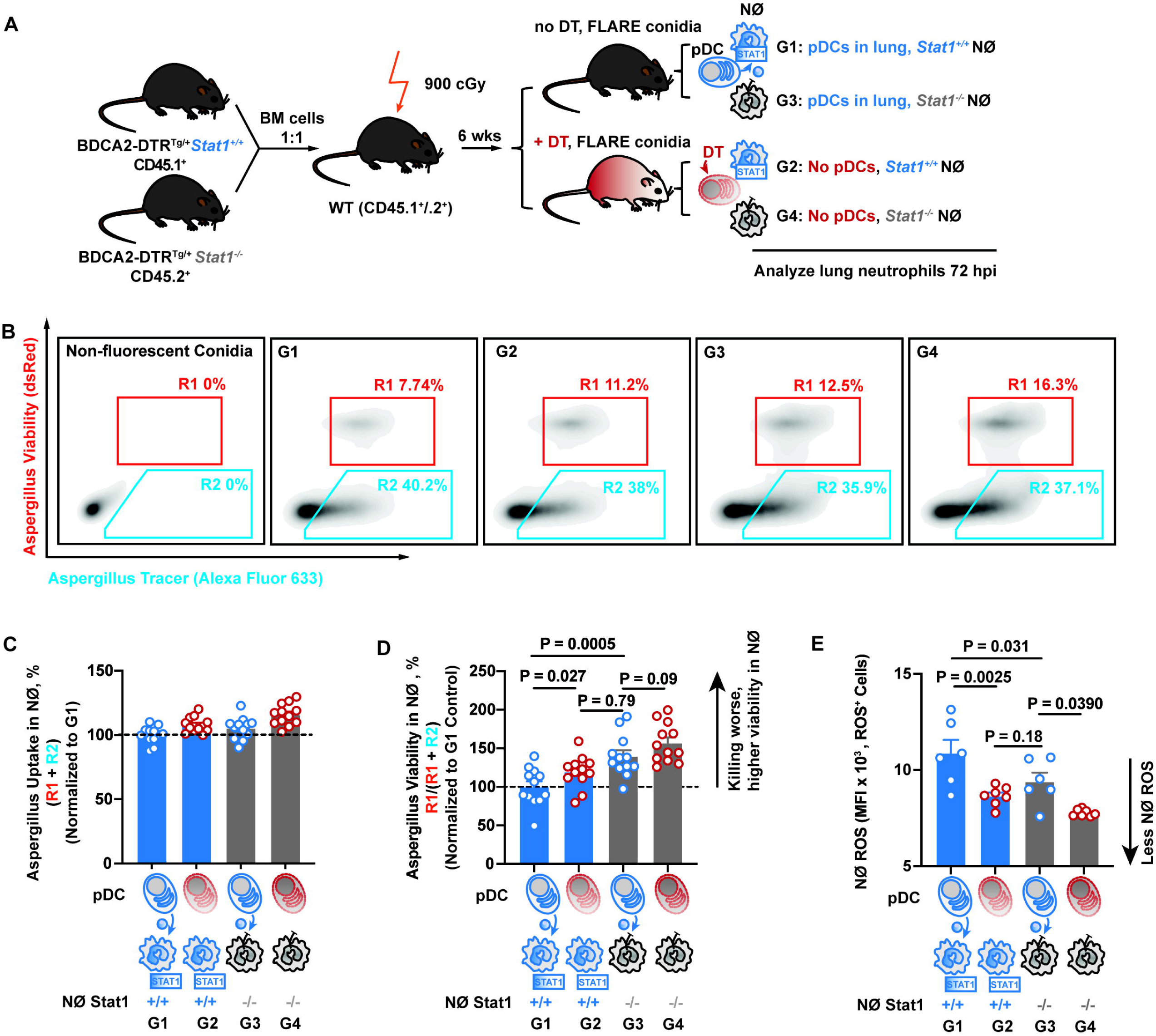
pDCs regulate neutrophil STAT1-dependent antifungal activity. (A) Scheme to generate and to deplete pDCs or not in CD45.1^+^ BDCA2-DTR^Tg/+^ *Stat1^+/+^* and CD45.2^+^ BDCA2-DTR^Tg/+^ *Stat1^-/-^* mixed bone marrow chimeric mice, resulting in 4 experimental groups (G1-G4). (B-D) (B) Representative plots that display DsRed and Af633 fluorescence intensity in lung neutrophils from 4 experimental groups: G1: pDC^+^ *Stat1^+/+^*(*Stat1^+/+^* neutrophils from pDC-sufficient mice), G2: pDC^-^ *Stat1^+/+^* (*Stat1^+/+^*neutrophils from pDC-depleted mice), G3: pDC^+^ *Stat1^-/-^* (*Stat1^-/-^* neutrophils from pDC-sufficient mice), G4: pDC^-^ *Stat1^-/-^* (*Stat1^-/-^*neutrophils from pDC-depleted mice). The R1 gate indicates the frequency of neutrophils that contain live conidia, the gate R2 indicates the frequency of neutrophils that contain killed conidia. (C and D) The plots show mean neutrophil (C) conidial uptake (R1 + R2) ± SEM and (D) conidial viability (R1/ (R1 + R2) ± SEM in indicated lung neutrophils isolated from the 4 groups, n = 12 per group, data pooled from 2 experiments. (E) Mean ± SEM ROS production in indicated lung neutrophils, n = 6 per group. (B - D) Infection dose: 3 × 10^7^ Af293 FLARE conidia via intratracheal route, analysis 72 hpi. (C and D) Data were pooled from 2 independent experiments and presented as mean ± SEM, (E) Data are representative of 2 experiments. Dots represent individual mice. Statistical analysis: Kruskal-Wallis test. See also Figure S3.

This experimental approach yielded 4 groups of neutrophils that were analyzed 72 hpi with FLARE conidia. Group 1 (G1) neutrophils were *Stat1^+/+^*neutrophils isolated from pDC-sufficient mice; G2 neutrophils were *Stat1^+/+^* neutrophils isolated from pDC-ablated mice; G3 neutrophils were *Stat1^-/-^* neutrophils isolated from pDC-sufficient mice; and G4 neutrophils were *Stat1^-/-^* neutrophils isolated from pDC-ablated mice (Fig. 4A). There was no difference in conidial uptake among these four groups of neutrophils (Fig. 4B and 4C), indicating that pDCs do not control neutrophil conidial uptake and that neutrophil-intrinsic STAT1 signaling is dispensable for this process.

The frequency of neutrophils that contained live conidia was markedly increased in G3 (pDC^+^ lungs; *Stat1^-/-^*) neutrophils compared to G1 (pDC^+^ lungs; *Stat1^+/+^*) neutrophils (Fig. 4B and 4D), consistent with prior experimental results (Fig. 3E). Critically, pDC ablation increased the frequency of *Stat1*^+/+^ neutrophils that contained live conidia (comparison of G2 versus G1 neutrophils; P = 0.027, Fig. 4B and 4D). This result indicates that pDC-derived products contribute to STAT1-dependent neutrophil conidiacidal activity. In contrast, pDC ablation did not significantly increase the frequency of *Stat1*^-/-^ neutrophils that contained live conidia (comparison of G4 versus G3 neutrophils; P = 0.09, Fig. 4B and 4D).

To obtain a complementary measurement of pDC to neutrophil STAT1 crosstalk, we measured ROS production in the 4 neutrophil groups in parallel. Consistent with the direct measurements of fungicidal activity in Fig. 4D and with the ROS measurements in Fig. 3I and 3J, *Stat1^+/+^* neutrophils displayed a higher ROS mean fluorescence intensity (MFI) than *Stat1*^-/-^ neutrophils (G1 versus G3). pDC ablation markedly reduced ROS production by *Stat1*^+/+^ neutrophils (G2 versus G1), in line with the reduction in neutrophil fungicidal activity (Fig. 4E). In fact, *Stat1*^+/+^ neutrophils isolated from pDC-depleted lungs (G2) had a similar ROS MFI as *Stat1*^-/-^ neutrophils isolated from pDC-sufficient lungs (G3). ROS levels observed in *Stat1*^-/-^ neutrophils isolated from pDC-depleted lungs (G3) was lower than the ROS levels observed in *Stat1*^-/-^ neutrophils isolated from pDC-sufficient lungs (Fig. 4E), consistent with the idea that pDCs can further modulate neutrophil ROS production in a *Stat1*-independent fashion. Collectively, these findings indicate that pDCs regulate STAT1-dependent neutrophil fungicidal activity and ROS production.

### STAT1-dependent guanylate-binding proteins are dispensable for the neutrophil anti-fungal activity

To gain further insight into how STAT1 regulates cell-intrinsic neutrophil fungicidal activity and ROS generation during respiratory *A. fumigatus* challenge, we generated mixed bone marrow chimeric mice (1:1 mix of CD45.2^+^ *Stat1*^−/−^ and CD45.1^+^ *Stat1*^+/+^ donor bone marrow cells → lethally irradiated CD45.1^+^CD45.2^+^ *Stat1*^+/+^ recipient mice) and performed bulk RNA-seq on *Stat1*^−/−^ and *Stat1*^+/+^ neutrophils sorted from *A. fumigatus*-infected recipient mice. There were marked differences in the transcriptome of *Stat1*^−/−^ and *Stat1*^+/+^ lung neutrophils (Fig. 5A), with 2586 genes showing differential expression in *Stat1*^−/−^ neutrophils compared to *Stat1*^+/+^ neutrophils. KEGG pathway enrichment analysis showed downregulation of many pathways in *Stat1*^−/−^ neutrophils, including the cytosolic DNA sensing pathway, proteasome, RIG-I like receptor signaling pathway, Toll-like receptor signaling pathway, cytokine-cytokine receptor pathway (Fig. 5B). We identified commonly downregulated genes in *Stat1*^−/−^ neutrophils, many of which were known interferon-regulated genes (ISGs; Fig. 5C).

**Figure 5.**
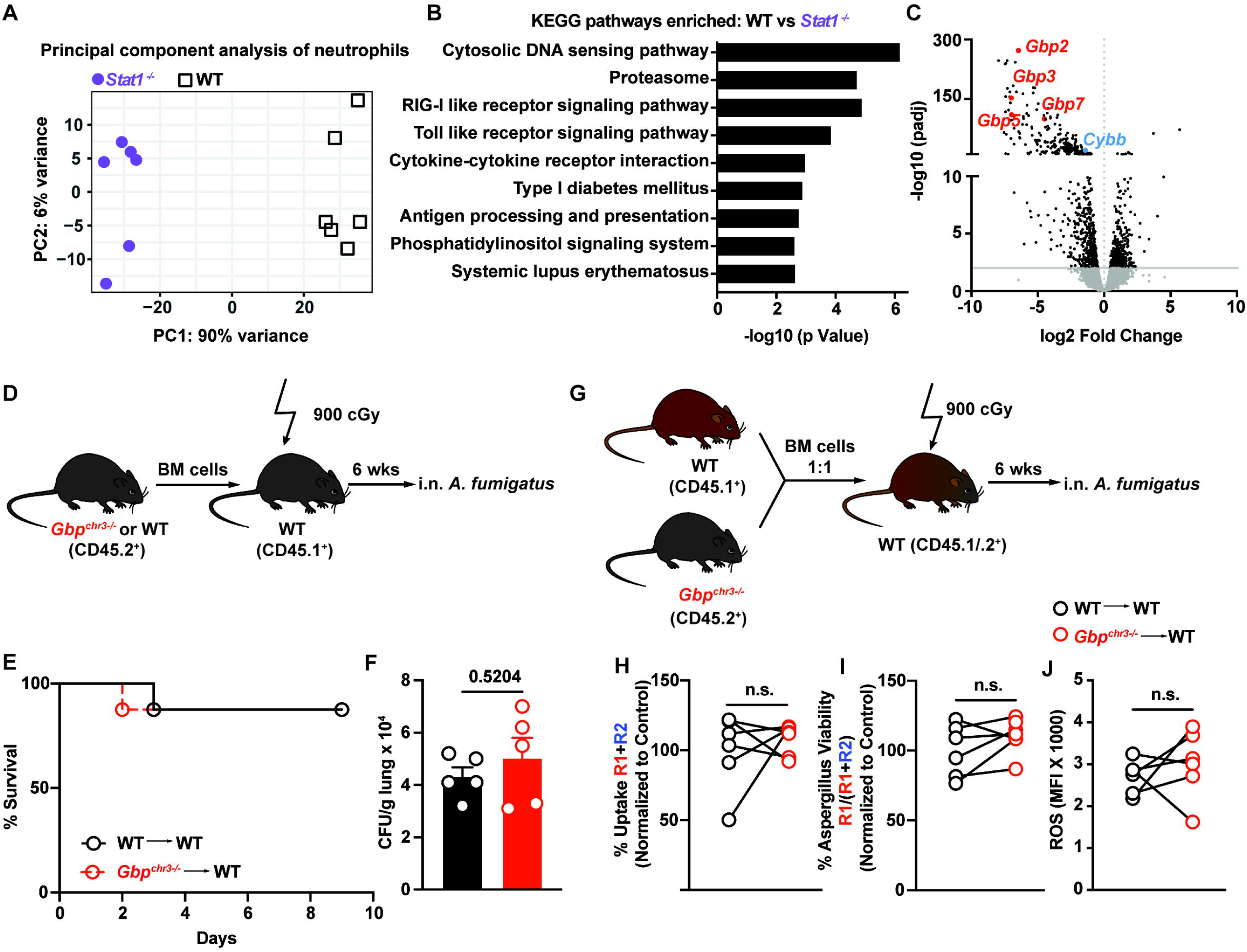
STAT1-dependent guanylate-binding proteins are dispensable for the neutrophil antifungal response. (A) Principal component analysis of global gene expression, (B) differentially enriched KEGG pathways, and (C) Volcano plot of differentially expressed genes in *Stat1^-/-^* (purple symbols) and *Stat1^+/+^*(black symbols) neutrophil sorted from mixed bone marrow chimeric mice (n = 6) at 72 hpi with 3 × 10^7^ CEA10 conidia. Selected downregulated genes in *Stat1^-/-^* neutrophils are highlight in the Volcano plot. (D) Experimental scheme to generate *GBP^chr3-/-^*and *GBP^chr3+/+^* single chimeric mice. (E) Kaplan Meier survival (n = 7-8 per group), and (F) mean ± SEM lung CFU (n = 5 per group) in *GBP^chr3-/-^* (red symbols) and *GBP^chr3+/+^* (black symbols) single chimeric mice infected with 3-6 × 10^7^ CEA10 conidia. (G) Experimental scheme to generate *GBP^chr3-/-^* and *GBP^chr3+/+^* mixed chimeric mice. (H and I) The plots show normalized neutrophil (H) conidial uptake (R1 + R2) ± SEM and (I) conidial viability (R1/ (R1 + R2) ± SEM in lung neutrophils isolated from *GBP^chr3-/-^* (red symbols) and *GBP^chr3+/+^* (black symbols) mixed bone marrow chimeric mice (n = 6 per group). (J) Mean ± SEM neutrophil ROS production in neutrophils isolated from *GBP^chr3-/-^* (red symbols) and *GBP^chr3+/+^* (black symbols) mixed bone marrow chimeric mice (n=6 per group). (H - J) Infection dose: 3 × 10^7^ Af293 FLARE conidia via intratracheal route, analysis 72 hpi. (F, H - J) Data are representative of 2 experiments and presented as mean ± SEM. Dots represent individual mice. Statistical analysis: Mann-Whitney test.

Several GTPases guanylate-binding proteins (GBPs), including *Gbp2*, *Gbp3*, *Gbp5* and *Gbp7* were significantly downregulated in *Stat1*^−/−^ neutrophils (Fig. 5C). To address the function of these genes in neutrophil antifungal activity in otherwise immune competent mice, we utilized Gbp^chr3−/−^ mice, which lack the entire chromosome 3 cluster that contains *Gbp1*, *Gbp2*, *Gbp3*, and *Gbp5*. We generated single chimeric mice (CD45.2^+^ Gbp^chr3−/−^ or CD45.2^+^ Gbp^chr3+/+^ → lethally irradiated CD45.1^+^ Gbp^chr3+/+^ recipients) (Fig. 5D) and compared the mortality and fungal burden of Gbp^chr3−/−^ and Gbp^chr3+/+^ chimeric mice. As expected from a prior study in corticosteroid and cyclophosphamide-treated mice (21), there was no difference in mortality or fungal colony forming unit (CFU) between Gbp^chr3−/−^ and Gbp^chr3+/+^ chimeric mice (Fig. 5E and 5F). Using mixed chimeric mice (CD45.2^+^ Gbp^chr3−/−^ and CD45.1^+^ Gbp^chr3+/+^ → CD45.1^+^CD45.2^+^ Gbp^chr3+/+^ recipients) (Fig. 5G) and FLARE conidia, we quantified the cell-intrinsic antifungal activity of Gbp^chr3−/−^ and Gbp^chr3+/+^ leukocytes and found no difference in neutrophil conidial uptake and killing (Fig. 5H and 5I). Moreover, Gbp^chr3−/−^ and Gbp^chr3+/+^ neutrophils isolated from the same lung exhibited no difference in ROS MFI (Fig. 5J).

The pDC-IFN-STAT1 axis regulates neutrophil *Cybb* expression during A*. fumigatus* infection.

Since GBPs did not contribute to STAT1-regulated neutrophil defense against *A. fumigatus*, we investigated other candidate genes that were downregulated in *Stat1*^−/−^ neutrophils (Fig. 5C) and focused next on *Cybb,* which encodes CYBB, the p91 subunit of the NADPH oxidase complex (NOX2). The four genes (*Cyba*, *Ncf1*, *Ncf2*, and *Ncf4*) that encode other subunits of the neutrophil NADPH oxidase complex (CYBA/p22 subunit, NCF1/p47 subunit, NCF2/p67 subunit and NCF4/p40 subunit) were not downregulated in *Stat1*^−/−^ neutrophils (Fig. 6A). During *A. fumigatus* infection lung *Cybb* expression peaked at 48 and 72 hpi (Fig. S4A), temporally coincident with the observed peaks in lung pDC influx and type I and type III IFN expression (Fig. 2A-C). The production of reactive oxygen species (ROS) by neutrophils is a vital mechanism for effectively eradicating fungal infections. Consistent with a prior study, we confirmed that *Cybb*-deficient mice (*gp91^phox-/-^*) were highly susceptible to *A. fumigatus* infection (Fig. S4B) (22). To investigate the hypothesis that STAT1 signaling in neutrophils regulates *Cybb* expression, we isolated neutrophils from infected *Stat1*^−/−^ and *Stat1*^+/+^ mice and found that *Cybb* mRNA level were decreased in *Stat1*^−/−^ neutrophils compare to *Stat1*^+/+^ neutrophils (Fig. 6B), consistent with STAT1-dependent regulation of *Cybb* transcription.

**Figure 6.**
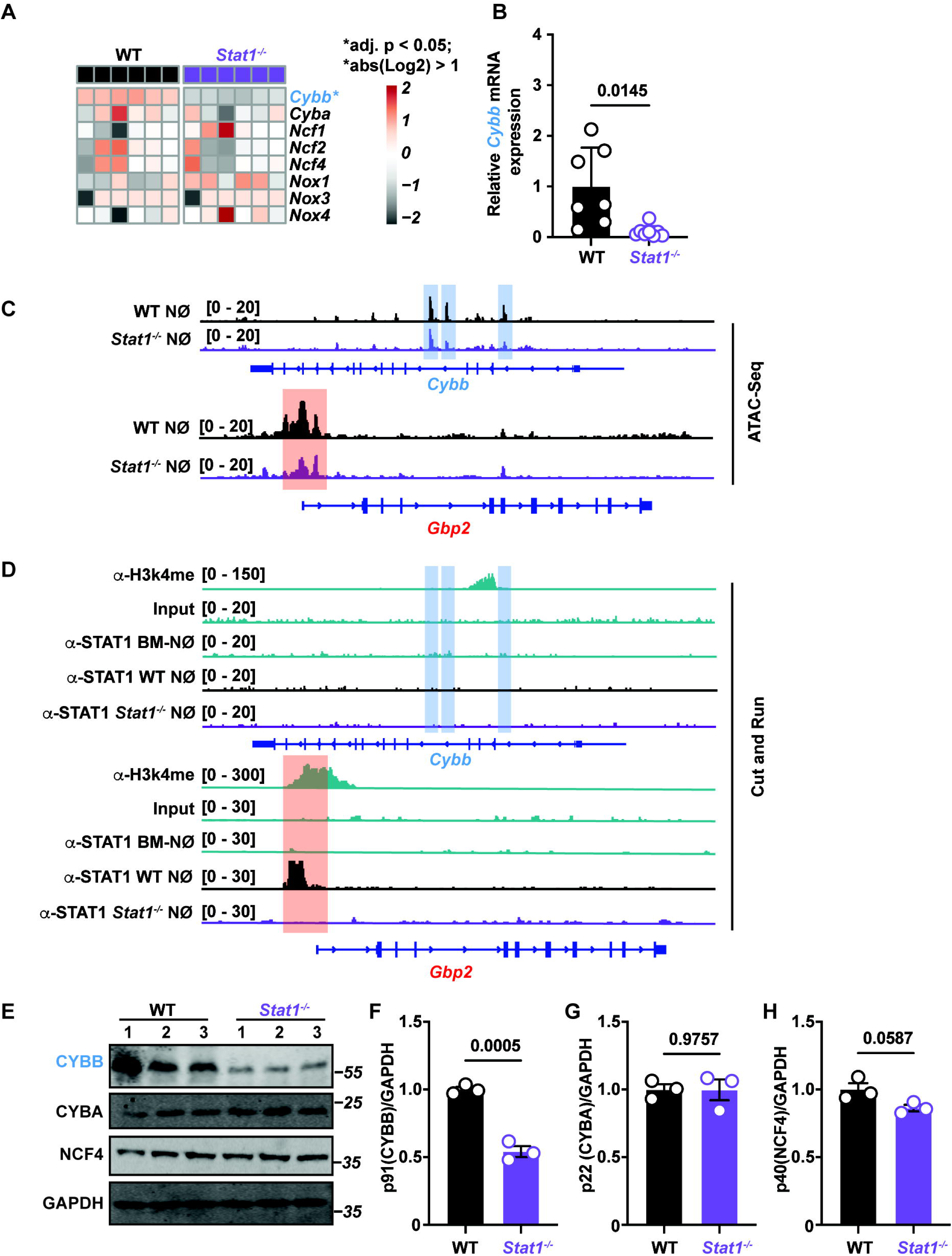
STAT1-dependent control of *Cybb* expression and CYBB protein levels in neutrophils. **(A)** Heatmap of *Cybb*, *Cyba*, *Ncf1*, *Ncf3*, *Ncf4*, *Nox1*, *Nox3*, and *Nox4* expression in neutrophils sorted from *Stat1^-/-^*(purple symbols) and *Stat1^+/+^*(black symbols) mixed bone marrow chimeric mice (n = 6). Genes with a STAR* are DEG. Each lane is an independent biological replicate. (B) qRT-PCR of *Cybb* mRNA expression in neutrophils sorted from *Stat1^-/-^* (purple symbols) and *Stat1^+/+^* (black symbols) mice. (C) ATAC-seq analysis of neutrophils were sorted from *Stat1^-/-^* (purple peaks) and *Stat^+/+^* (black peaks) mice. Gene tracks show chromatin accessibility at the *Cybb* and *Gbp2* locus. (D) Bone marrow and lung neutrophils were sorted from *Stat1^-/-^*(purple peaks) and *Stat1^+/+^* (black peaks) mice and processed for CUT&RUN. Gene tracks show STAT1 signal as normalized fragment pileup (y-axis) plotted by genome position (x-axis). The shaded box highlights a putative STAT1 binding site at the Gbp2 promoter region. Gene tracks show H3K4me3 ChIP-seq signal as normalized fragment pileup (top rows; green). (E - H) Western blot and quantitation of (F) CYBB, (G) CYBA, (H) NCF4 vs GAPDH protein levels in lung neutrophils sorted from *Stat1^-/-^* (purple symbols) and *Stat1^+/+^* (black symbols) mice, neutrophils from 4-5 mice were pooled to obtain sufficient protein for Western blot analysis in each biological replicate. (A - H) Infection dose: 3 × 10^7^ CEA10 conidia via intratracheal route, analysis 72 h pi. (C) Data were calculated by Kruskal-Wallis nonparametric test for multiple comparisons for each group compared with control group. (G – I) Data are representative of 2 experiments. Dots represent individual mice. Statistical analysis: Mann-Whitney test. See also Figure S4.

To determine chromatin accessibility at the *Cybb* locus, we performed assay for transposase-accessible chromatin (ATAC) sequencing of sorted *Stat1*^−/−^ and *Stat1*^+/+^ lung neutrophils from *A. fumigatus*-infected mice. Within the *Cybb* locus, we found three regions (blue squares) that were less accessible in *Stat1*^−/−^ neutrophils than in *Stat1*^+/+^ neutrophils (Fig. 6C), consistent with a modest STAT1-dependent regulation of *Cybb* expression (Fig. 6A and 6B). As a positive control, within Gbp2 locus, we found one region (red square) that was less accessible in *Stat1*^−/−^ neutrophils compared to *Stat1*^+/+^ neutrophils (Fig. 6C), consistent with STAT1-dependent regulation of *Gbp* expression (Fig. 5C).

To probe STAT1 binding to the Cybb gene locus, we performed <u>C</u>leavage <u>U</u>nder <u>T</u>argets <u>&</u> <u>R</u>elease <u>U</u>sing <u>N</u>uclease (CUT&RUN) experiments of sorted *Stat1*^−/−^ and *Stat1*^+/+^ lung neutrophils from *A. fumigatus*-infected mice. We did not observe direct STAT1 binding to *Cybb* locus and promoter region (Fig. 6D). In CUT&RUN experiments, STAT1 bound to the *Gbp2* locus and promoter region (Fig. 6D). Collectively, these results suggest that STAT1 regulates *Cybb* transcription via an indirect mechanism.

Because STAT1 did not appear to regulate *Cybb* transcription through direct effects on chromosomal accessibility or binding to the *Cybb* locus, we next explored whether lung neutrophil CYBB protein levels were regulated by STAT1 signaling. *Stat1*^−/−^ and *Stat1*^+/+^ neutrophils were isolated from *A. fumigatus* infected mice lungs at 72 hpi and analyzed for CYBB expression by Western blotting. CYBB protein levels, but not other subunits (p22/CYBA and p40/NCF4) of NADPH oxidase protein levels were lower in *Stat1*^−/−^ neutrophils compared to *Stat1*^+/+^ neutrophils (Fig. 6E – 6H), linking neutrophil STAT1 activation to CYBB expression, and to the oxidative burst.

## Discussion

Our data introduce a model of protective pDC to neutrophil crosstalk in which pDCs undergo a defined activation trajectory in transit to the *Aspergillus*-infected lung. pDC activation provides a critical source of type I and type III IFN at the portal of infection, and licenses neutrophil antifungal properties in the lung in a direct manner. The latter step occurs through neutrophil-intrinsic STAT1-dependent control of CYBB protein levels. Local pDC-dependent control of NADPH oxidase assembly regulates the strength of neutrophil oxidative burst, as judged by reactive oxygen species production, boosts neutrophil intracellular conidial killing, and confers sterilizing immunity against inhaled spores.

pDCs originate in the bone marrow (BM), travel through the bloodstream, and migrate to lymphoid and nonlymphoid tissues during both normal and inflammatory states (18, 23–25). In a recent study we demonstrated that *Aspergillus*-infected Mo-DCs and neutrophils release CXCR3 ligands (i.e., CXCL9 and CXCL10) into the inflamed lung, coupling fungal recognition and fungus-induced inflammation to CXCR3 signaling-dependent pDC influx from the circulation into the lung. In this study, *A. fumigatus* infection induces a pDC activation trajectory that substantially alters the lung rather than the BM pDC transcriptome, implying that pDC trafficking to peripheral tissue precedes activation occurs at the portal of infection. While not a focus of this work, the precise mechanism of pDC activation in the fungus-infected lung remains an open question, in part because conditional gene deletion strategies do not exist for pDCs, precluding facile comparison of gene-deficient and gene-sufficient pDCs in infected tissues in mice with no other genetic perturbations.

Our data indicate that lung-infiltrating pDCs upregulate genes implicated in fungal recognition, including the C-type lectin receptors (CLRs) and downstream signaling molecules. pDCs express the C-type lectin receptors Dectin1, Dectin2, and Dectin3 and can secrete IFN*α* and TNF when co-incubated with *Aspergillus* hyphae via Dectin2 signaling (26–28), though the interaction of pDCs with conidia, the infectious propagules, were not examined. pDCs do not internalize *Aspergillus* conidia readily in the test tube or in the infected lung (4, 26), supporting the notion that activation occurs via fungal cell contact or via the presence of activating cytokines or other inflammatory mediators. Support for the latter scenario comes from an experiment in which curdlan (i.e., a particulate Dectin1 agonist)-induced upregulation of pDC CD40 and CD83 expression could be increased by simultaneous co-incubation with an acellular curdlan-stimulated peripheral blood mononuclear cell supernatant (26). Beyond *Aspergillus*, pDC-enriched human cell fractions can release TNF in a Dectin2 and Dectin3 signaling-dependent fashion in response to *Paracoccidioides brasiliensis* co-incubation (29). Beyond CLR signaling, we found that *Aspergillus* infection triggered upregulation of Toll-like receptors in lung-infiltrating pDCs. Prior work demonstrated that *Aspergillus*-derived unmethylated CpG sequences can activate TLR9 signaling *in vitro* (30, 31), and this may represent an additional mechanism by which pDCs become activated *during Aspergillus* infection.

During Dengue, Zika, and Hepatitis C viral infections, physical contact with virally infected cells stimulates pDC-mediated antiviral responses (32, 33). α_L_β_2_ integrin (lymphocyte function-associated antigen-1; LFA1) expressed by the pDC can bind to ICAM1 on infected cells to promote a sustained interaction, termed an interferonogenic synapse, during which viral RNA is transferred to pDCs, leading to IFN production via the nucleic acid sensor TLR7. This process activates type I IFN-dependent antiviral programs in infected tissues. pDC commitment to type I IFN production is further regulated by antecedent cell-intrinsic TNF receptor and leukemia inhibitory factor signaling during murine cytomegalovirus infection, underscoring the contribution of local cytokine signaling to pDC activation by contact-dependent and - independent mechanisms (34). The lung pDC transcriptomic data detected increased expressed ICAM1 in the lung-infiltrating pDCs isolated from infected mice. This observation raises the possibility that ICAM1 expression may facilitate lung pDC contact with β_2_-integrin-expressing myeloid cells to facilitate reciprocal interactions. In sum, the upregulation of multiple receptors from various classes suggests that pDCs likely employ a blend of receptors to identify and react to fungal pathogens, leading to full pDC activation in the fungus-infected lung. Notably, we were unable to isolate sufficient pDCs to compare the bulk lung pDC transcriptome in naïve mice with the lung pDC transcriptome in *Aspergillus*-infected mice.

Recent studies have advanced the concept that circulating neutrophils exhibit transcriptional heterogeneity in the steady state and during microbial infection (35). In addition, neutrophils can acquire transcription-dependent non-canonical functions upon entry into peripheral tissues, exemplified by a regulatory role in angiogenesis in the lung (36). Single-cell analysis has revealed the presence of three separate circulating murine and human populations that differ with respect to interferon-stimulated gene (ISG) and CXCR4 expression. Interestingly, all three populations expressed similar levels of *Cybb* mRNA and exhibited similar NADPH oxidase scores by gene ontogeny analysis during homeostasis and systemic bacterial (i.e., *Escherichia coli*) infection (35). During tuberculosis, malaria, and hematopoietic cell transplantation, circulating human neutrophils exhibit an IFN transcriptional signature (37–39), which in the case of malaria, is reduced by receipt of anti-malarial therapy. The finding that pDC-derived type I and type III IFN licenses neutrophil antifungal activity through STAT1-dependent control of *Cybb* transcription and CYBB protein levels raises the important question whether the ISG^hi^ neutrophil subset is enriched in the lung and particularly effective at killing *Aspergillus* conidia compared to the other two ISG^lo^ circulating subsets. The ability to track and quantify fungal uptake killing with FLARE conidia will facilitate future studies to link lung-infiltrating neutrophil subsets, defined by distinct transcriptional profiles, to the quality of effector functions at single-encounter resolution. These studies support idea that dynamic transcriptional plasticity represents a cardinal feature of the circulating neutrophil response to microbial infection and other external stressors. It remains unclear whether the observed transcriptional plasticity represents dynamic changes in the production of transcriptionally heterogeneous circulating neutrophil subsets or represents an interconversion between circulating subsets.

The effects of type III IFN on neutrophils are context-dependent. In a DSS model of sterile colitis, IFNλ has been linked to neutrophil ROS suppression and a reduction in neutrophil degranulation (40). In human neutrophils, recombinant IFNλ can inhibit platelet-induced NETosis (41). IFNλ has various roles in bacterial infections (42, 43), with a previous study demonstrated that *Bordetella pertussis* infection induces IFNλ and IFNλ receptor (IFNLR1) expression as well as inflammation in the lung (42). IFNλ signaling in neutrophils can suppress *B. pertussis* killing and neutrophil production of ROS, MMP9, NETs, MPO, and IFNγ (44, 45). In contrast, conditional deletion of the IFNλ receptor in neutrophils is linked to a reduction in neutrophil ROS during pulmonary *A. fumigatus* infection (15). During HSV corneal infection, application of recombinant IFNλ suppressed neutrophil recruitment but did not impact virucidal activity or ROS production (46). These results indicate that IFNλ may have different biological effects based on cellular targets and responsiveness at sites of inflammation and on the type of inflammatory stimulus. In this study, the STAT1-dependent action of pDC-dependent IFNs (both type I and III) on neutrophils is critical for their local licensing of their cell-intrinsic cytotoxic activity against *Aspergillus* conidia. The individual contribution of type I versus type III IFN-dependent activation on STAT1-dependent CYBB expression remains unclear. An open question remains how STAT1 signaling precisely regulates *Cybb* transcription and CYBB translation, since we could not detect clear evidence of STAT1 binding to the *Cybb* promoter or clear STAT1-dependent changes in chromatin accessibility.

Overall, these findings provide important insights into the pDC-STAT1-CYBB axis as a key regulator of NADPH oxidase expression and highlight the critical role of this pathway in promoting antifungal immunity in neutrophils. The discovery that pDCs can regulate ROS induction by neutrophils by controlling the STAT1-dependent expression of a single NADPH oxidase subunit adds to our understanding of the complex and protective interplay between innate immune cells during fungal infection.

## Supporting information

Related to Fig. 1

Related to Fig. 3

Related to Fig. 4

Related to Fig. 6

## ACKNOWLEDGMENTS

We thank Dr. Thirumala-Devi Kanneganti (St. Jude Children’s Research Hospital) providing the *GBP^chr3-/-^* bone marrows. We thank the MSKCC IGO core for performing RNA sequencing, and Chen Liao (Geisel School of Medicine at Dartmouth), Paul Zumbo (Weill Cornell Medicine) for assistance with the RNA sequencing data analysis. We thank Robert Cramer (Geisel School of Medicine at Dartmouth), Iliyan Iliev (Weill Cornell Medicine), and members of the Hohl laboratory for numerous conversations and insights into this study. In this study, T. M. H. was supported by NIH grants P30 CA 008748 (to MSKCC), R37 AI093808, and R01 AI139632; Y. G. was supported by the Ludwig Center Post-Doctoral Award, MSKCC; M.A.A. was supported by NIH F31 AI161996; K.A.M.M. was supported by NIH F31 AI167511; K. B. M. was supported by Experimental Immuno-oncology Scholars’ Program, MSKCC and NIH 5T32CA009149; J. C. S. was supported by the Ludwig Center for Cancer Immunotherapy, the American Cancer Society, the Burroughs Wellcome Fund, and the NIH grants R01 AI100874, R01 AI130043, R01 AI155558, and P30CA008748. S. G. was supported by CRI / Donald J. Gogel Postdoctoral Fellowship CRI Award CRI3934 and NIH K99AI180360. The funders had no role in study design, data collection and analysis, decision to publish or preparation of manuscript. We acknowledge the use of the Integrated Genomics Operation Core, funded by the NCI Cancer Center Support Grant (CCSG, P30 CA08748), Cycle for Survival, and the Marie-Josée and Henry R. Kravis Center for Molecular Oncology.

## AUTHOR CONTRIBUTIONS

Conceptualization, T.M.H., and Y.G.; Methodology, T.M.H., and Y.G.; Investigation, Y.G., M.A.A., K.A.M.M, S. A. G., H. K., P. Z., M.G., A.B., K.M., Y. Y.; Writing – Original Draft, Y.G. and T.M.H.; Writing – Review & Editing, Y.G. and T.M.H.; Funding Acquisition, T.M.H. and Y.G.; Resources, J. S., and D. B.

## MATERIALS AND METHODS

### Chemicals and reagents

Unless otherwise noted, chemicals were purchased from Sigma-Aldrich or Fisher Scientific, cell culture reagents from Thermo Fisher Scientific, and microbiological culture media from BD Biosciences. Antibodies for flow cytometry were acquired from BD Biosciences, Thermo Fisher Scientific and Tonbo.

### Mice

Our study examined male and female animals, and similar findings are reported for both sexes. C57BL/6J (JAX: 00664), BDCA2-DTR (JAX: 014176) mice were from Jackson Laboratories. *GBP^chr3-/-^* bone marrows were provided by Dr. Thirumala-Devi Kanneganti. C57BL/6.SJL (Stock: 4007) were purchased from Taconic. C57BL/6 and C57BL/6.SJL mice were crossed to generate CD45.1^+^CD45.2^+^ recipient mice for mixed BM chimeras. CD45.2^+^ BDCA2-DTR^Tg/+^ were backcrossed to C57BL/6.SJL mice to obtain CD45.1^+^ BDCA2-DTR^Tg/+^ mice. BDCA2-DTR^Tg/+^ were crossed with CD45. 2^+^ *Stat1*^-/-^ mice to obtain CD45. 2^+^ BDCA2-DTR^Tg/+^ *Stat1*^-/-^ mice. For experiments in which the breeding strategy did not yield littermate controls, gene-knockout mice were co-housed with C57BL/6 mice for 14 days prior to infection, whenever possible.

### Generation of Bone Marrow Chimeric Mice

For single BM chimeras, CD45.1^+^ C57BL/6.SJL recipients were lethally irradiated (900cGy), reconstituted with either 2-5 × 10^6^ CD45.2^+^ *GBP^chr3-/-^*, CD45.2^+^ C57BL/6J or CD45.2^+^ *Stat1*^-/-^ BM cells. For mixed BM chimeras, CD45.1^+^CD45.2^+^ recipients were irradiated and reconstituted with a 1:1 mixture of CD45.1^+^ C57BL/6.SJL and CD45.2^+^ *Stat1*^-/-^ or CD45.2^+^ *GBP^chr3-/-^* BM cells, CD45.1^+^ BDCA2-DTR^Tg/+^ and CD45.2^+^ BDCA2-DTR^Tg/+^ *Stat1*^-/-^. After BM transplantation, recipient mice received 400 μg/ml enrofloxacin in the drinking water for 21 days to prevent bacterial infections and rested for 6-8 weeks prior to experimental use.

### Aspergillus fumigatus Infection Model

*A. fumigatus* Af293, Af293-dsRed (20), and CEA10 (47) strains were cultured on glucose minimal medium slants at 37°C for 4-7 days prior to harvesting conidia for experimental use. To generate FLARE conidia, briefly, 7 × 10^8^ Af293-dsRed conidia were rotated in 10 μg/ml Biotin XX, SSE in 1 ml of 50 mM carbonate buffer (pH 8.3) for 2 hr at 4 °C, incubated with 20 μg/ml Alexa Fluor 633 succinimidyl ester at 37 °C for 1 h, resuspended in PBS and 0.025% Tween 20 for use within 24 hrs. (20, 48). To generate morphologically uniform heat-killed swollen conidia, 5X10^6^/ml conidia were incubated at 37° C for 14 hours in RPMI-1640 and 0.5 μg/ml voriconazole and heat killed at 100 °C for 30 minutes (49). To infect mice with 30-60 million live or heat-killed *A. fumigatus* cells, conidia were resuspended in PBS, 0.025% Tween-20 at a concentration of 0.6-1.2X10^9^ cells and 50 μl of cell suspension was administered via the intranasal route to mice anesthetized by isoflurane inhalation.

### Analysis of *in vivo* and *in vitro* conidial uptake and killing

To analyze of conidia uptake and killing, FLARE conidia were used to infect mice. In data analyses for a given leukocyte subset, conidial uptake refers to the frequency of fungus-engaged leukocytes, i.e., the sum of dsRed^+^ AF633^+^ and dsRed^-^ AF633^+^ leukocytes. Conidial viability within a specific leukocyte subset refers to the frequency of leukocytes that contain live conidia (dsRed^+^ AF633^+^) divided by the frequency of all fungus-engaged leukocytes (dsRed^+^ AF633^+^ and dsRed^-^ AF633^+^).

### *In vivo* Cell Depletion

To ablate specific cells, BDCA2-DTR^Tg/+^, BDCA2-DTR^Tg/+^ *Stat1*^-/-^, and non-transgenic littermate controls were injected i.p. with 10 ng/g body weight DT on Day -1, Day 0, and Day +2 pi (19), unless noted otherwise.

### Analysis of Infected Mice

To prepare single cell lung suspensions for flow cytometry, we followed the method outlined in (Hohl et al., 2009) with minor modifications. After perfusing murine lungs, they were placed in a gentle MACS™ C tube and mechanically homogenized in 5 ml RPMI-1640, 10% FBS, and 0.1 mg/ml DNAse using a gentle MACS™ Octo Dissociator (Miltenyi Biotech) without the use of collagenase. The lung cell suspensions were then lysed of RBCs, enumerated, and stained with fluorophore-conjugated antibodies prior to flow cytometric analysis. We used either a Cytoflex or a BD Aria for flow cytometric sorting and performed flow plot analysis using FlowJo v.9.6.6 software.

Neutrophils were identified as CD45^+^ CD11b^+^ Ly6C^lo^ Ly6G^+^ cells, monocytes as CD45^+^ CD11b^+^ CD11c^-^ Ly6G^-^ Ly6C^hi^ cells, Mo-DCs as CD45^+^ CD11b^+^ CD11c^+^ Ly6G^-^ Ly6C^hi^ MHC class II^+^ cells, and pDCs as CD45^+^ CD11c^int^ SiglecF^-^ CD19^-^ NK1.1^-^ CD11b^-^ B220^+^ SiglecH^+^ cells.

To assess the lung fungal burden, perfused murine lungs were homogenized using a PowerGen 125 homogenizer (Fisher) in 2 mL of PBS containing 0.025% Tween-20 and plated onto Sabouraud dextrose agar. For analysis of cytokine levels by ELISA, whole lungs were weighed and mechanically homogenized in 2 mL of PBS containing protease inhibitors. To analyze cytokine levels by qRT-PCR, we extracted total RNA from cells using TRIzol (Invitrogen). One to two micrograms of total RNA were reverse-transcribed using the High-Capacity cDNA Reverse Transcription Kit (Applied Biosystems). We used TaqMan Fast Universal Master Mix (2×) and TaqMan probes (Applied Biosystems) for each gene and normalized to glyceraldehyde-3-phosphate dehydrogenase. Gene expression was calculated using the Ct method relative to the naïve sample.

Intracellular ROS levels were measured in cells using CM-H2DCFDA [5-(and 6-) chloromethyl-2,7-dichlorodihydrofluorescein diacetate, acetyl ester] as described in (50). Briefly, single cell lung suspensions were incubated with 1μM CM-H2DCFDA in Hanks’ balanced salt solution at 37° C for 45 min according to manufacturer’s instruction and analyzed by flow cytometry.

### Immunoblotting assay

Cells were lysed in lysis buffer (150 mM NaCl, 50 mM HEPES (pH 7.4), 1 mM EDTA, 1% Nonidet P-40, protease inhibitors). Total cell lysates were subjected to SDS-PAGE and then blotted using indicated antibodies, Cybb polyclonal antibody (Invitrogen, cat. no. PA5-76034), p22phox mAb (CST, cat. no. 37570), p40phox mAb (cat. no. AB76158), GAPDH mAb (CST, cat. no. 5174s).

### RNA sequencing

#### RNA extraction

Phase separation in cells lysed in 1 mL TRIzol Reagent (ThermoFisher cat. no. 15596018) was induced with 200 µL chloroform. RNA was extracted from 350 µL of the aqueous phase using the miRNeasy Micro or Mini Kit (Qiagen cat. no. 217084/217004) on the QIAcube Connect (Qiagen) according to the manufacturer’s protocol. Samples were eluted in 13-15 µL RNase-free water.

#### Transcriptome sequencing

After RiboGreen quantification and quality control by Agilent BioAnalyzer, 1.9-2.0 ng total RNA with RNA integrity numbers ranging from 7.8 to 10 underwent amplification using the SMART-Seq v4 Ultra Low Input RNA Kit (Clonetech cat. no. 63488), with 12 cycles of amplification. Subsequently, 7.4-10 ng of amplified cDNA was used to prepare libraries with the KAPA Hyper Prep Kit (Kapa Biosystems cat. no. KK8504) using 8 cycles of PCR. Samples were barcoded and run on a HiSeq 4000 or NovaSeq 6000 in a PE50 (HiSeq) or PE100 (NovaSeq) run, using the HiSeq 3000/4000 SBS Kit or NovaSeq 6000 S4 Reagent Kit (200 Cycles) (Illumina). An average of 40 million paired reads were generated per sample and the percent of mRNA bases per sample averaged 84%.

#### RNA-Sequencing data analysis

Raw reads were quality checked with FastQC v0.11.7 (http://www.bioinformatics.babraham.ac.uk/projects/fastqc/), and adapters were trimmed using Trim Galore v0.6.7 (http://www.bioinformatics.babraham.ac.uk/projects/trim_galore/). Reads were aligned to the mouse reference genome (GRCm38.p6) using STAR v2.6.0c (51) with default parameters. Gene abundances were calculated with featureCounts v1.6.2 (52) using composite gene models from Gencode release vM17. Principle component analysis was performed using the plotPCA function from DESeq2 v1.32.0 (53). Differentially expressed genes were determined with DESeq2 v1.32.0 with a two-factor model incorporating batch as a covariate, with significance determined by Wald tests (q < 0.05). Gene set enrichment analysis was performed using fgsea v1.18.0 (54) with gene sets from the Broad Institute’s MSigDB (55, 56) collections; genes were ranked by the DESeq2 Wald statistic. Only pathways with an adjusted P value < 0.05 were considered enriched. Expression heatmaps were generated using variance-stabilized data, with the values centered and scaled by row. Code has been deposited to GitHub (https://github.com/abcwcm/Guo2024).

### RNAscope microscopy

Formaldehyde-fixed, paraffin embedded (FFPE) surgical tissue Sects. FFPE tissue sections (5 µm) were processed exactly as described in the manufacturer’s instructions (ACD Bio, cat. no. 323100). Probes targeting the following genes were used: Ly6G (ACD Bio, cat. no. 455701-C3), IFN*λ*R (ACD Bio, cat. no. 512981-C1), IFN*α*R1(ACD Bio, cat. no. 512971-C2). Slides were mounted with Prolong Diamond mounting media (ThermoFisher Scientific, cat. no. P36965). Slides were scanned using a Pannoramic Digital Slide 517 Scanner (3DHISTECH, Budapest, Hungary) using a 20×/0.8NA objective.

### Assay for transposase-accessible chromatin (ATAC) sequencing and epigenome analyses

Freshly harvested WT and KO mouse neutrophils were directly sent to MSKCC’s Epigenetics Research Innovation Lab. ATAC was performed as previously described (Corces et al. Nature Methods 2017) using 50,000 cells per replicate and the Tagment DNA TDE1 Enzyme (Illumina, 20034198). Sequencing libraries were prepared using the ThruPLEX DNA-Seq Kit (Takarabio, R400676) and sent to the MSKCC Integrated Genomics Operation core facility for sequencing on a NovaSeq 6000. Raw sequencing reads were trimmed and filtered for quality (Q>15) and adapter content using version 0.4.5 of TrimGalore (https://www.bioinformatics.babraham.ac.uk/projects/trim_galore) and running version 1.15 of cutadapt and version 0.11.5 of FastQC. Version 2.3.4.1 of bowtie2 (http://bowtie-bio.sourceforge.net/bowtie2/index.shtml) was employed to align reads to mouse assembly mm10 and alignments were deduplicated using Mark Duplicates in Picard Tools v2.16.0. Enriched regions were discovered using MACS2 (https://github.com/taoliu/MACS) with a p-value setting of 0.001, filtered for blacklisted regions (http://mitra.stanford.edu/kundaje/akundaje/release/blacklists/mm10-mouse/mm10.blacklist.bed.gz), and a peak atlas was created using +/- 250 bp around peak summits. The BEDTools suite (http://bedtools.readthedocs.io) was used to create normalized bigwig files. Version 1.6.1 of featureCounts (http://subread.sourceforge.net) was used to build a raw counts matrix and DESeq2 was employed to calculate differential enrichment for all pairwise contrasts. Peak-gene associations were created by assigning all intragenic peaks to that gene, while intergenic peaks were assigned using linear genomic distance to transcription start sites (TSS).

### Histone and Transcription Factor CUT&RUN

For CUT&RUN, 400,000 sorted neutrophils were used for H3K4me3 and STAT1 analysis. Anti-H3K4me3 (Epicypher, cat. no. 13-0028) or polyclonal anti-STAT1 antibodies (Proteintech, cat. no. 10144-2-AP) were employed for each target. Sorted cells were washed with PBS and resuspended in Antibody Buffer (1 × eBioscience Perm/Wash Buffer, 1× Roche cOmplete EDTA-free Protease Inhibitor, 0.5 µM Spermidine, and 2 µM EDTA in H2O). They were incubated overnight at 4°C with control IgG, H3K4me3, or STAT1 antibodies diluted 1:100 in Antibody Buffer in a 96-well V-bottom plate. Following antibody incubation, cells were washed twice with Buffer 1 (1× eBioscience Perm/Wash Buffer, 1× Roche cOmplete EDTA-free Protease Inhibitor, 0.5 µM Spermidine in H2O) and resuspended in 50 µL of Buffer 1 plus 1× pA/G-MNase (Cell Signaling, cat. no. 57813). This mixture was incubated on ice for 1 hour, then washed twice with Buffer 2 (0.05% w/v Saponin, 1× Roche cOmplete EDTA-free Protease Inhibitor, 0.5 µM Spermidine in 1X PBS) three times. Cells were resuspended in Calcium Buffer (Buffer 2 plus 2 µM CaCl2) and incubated on ice for 30 minutes to activate the pA/G-MNase reaction. An equal volume of 2× STOP Buffer (Buffer 2 plus 20 µM EDTA plus 4 µM EGTA) and 1 pg of *Saccharomyces cerevisiae* spike-in DNA (Cell Signaling, cat. no. 29987) were added. Samples were incubated at 37 °C for 15 minutes, followed by DNA isolation and purification using the Qiagen MinElute Kit per the manufacturer’s instructions.

Immunoprecipitated DNA was quantified by PicoGreen, and the size was evaluated using an Agilent BioAnalyzer: fragments between 100 and 600 bp were size selected using aMPure XP beads (Beckman Coulter, cat. no. A63882). KAPA HTP Library Preparation Kit (Kapa Biosystems, cat. no. KK8234) was used to prepare Illumina sequencing libraries according to the manufacturer’s instructions with 0.001–0.5 ng input DNA and 14 cycles of PCR. Barcoded libraries were run on the NovaSeq 6000 in a PE100 run using S4 kit version 1.5 with XP mode to generate approximately 23 million paired reads per sample.

### CUT&RUN Data Processing

CUT&RUN datasets were processed by trimming paired reads for adaptors and low-quality sequences using Trimmomatic (v0.39) and aligning to the mm10 reference genome with Bowtie 2 (v2.4.1). Peaks were identified using MACS2 (v2.2.7.1) with input samples as control, employing narrow peak parameters and cutoff - analysis - ple-5 - keep - dup all -B - SPMR. Irreproducible discovery rate (IDR) calculations were performed using ENCODE project scripts (IDR v2.0.4.2). Reproducible peaks with an IDR value of 0.05 or less in each condition were retained, aggregated, and merged to create the final atlas, which was annotated with the UCSC Known Gene model. Reads were mapped to this atlas and counted using the summarize Overlaps function from the Genomic Alignment package (v1.34.1).

### Visualization

Genomic tracks were visualized using GViz (v1.42.1) or IGV (v2.9.4).

### Quantitation and Statistical Analysis

All data presented are representative of at least two independent experiments, as indicated. Unless stated otherwise, all results are expressed as mean (± SEM). We used the Mann-Whitney test for comparisons of two groups, and the Kruskal-Wallis test for multi-group comparisons, unless noted otherwise. Survival data were analyzed using the long-rank test. All statistical analyses were performed using GraphPad Prism software (v9.2.0).

### Study approval

All mouse strains were bred and housed in the MSKCC or St. Jude Children’s Research Hospital Animal Resource Center under specific pathogen-free conditions. All animal experiments were conducted with sex- and age-matched mice and performed with MSKCC Institutional Animal Care and Use Committee approval. Animal studies complied with all applicable provisions established by the Animal Welfare Act and the Public Health Services Policy on the Humane Care and Use of Laboratory Animals.

### Resource availability and Lead Contact

Further information and requests for resources or reagents should be directed to the Lead Contact, Tobias M. Hohl.

### Materials Availability

This study did not generate new unique reagents.

### Data and Code Availability

Values for all data points in graphs are reported in the Supporting Data Values file. RNA sequencing data, ATAC sequencing data and CUT&RUN sequencing data were uploaded into NCBI database. Raw RNA sequencing datasets generated in this study are available at GSE280164. Raw ATAC sequencing data and CUT&RUN sequencing data generated in this study are available at GSE280229.

### Key Resources Table

| REAGENT or RESOURCE | SOURCE | IDENTIFIER |
| --- | --- | --- |
| Antibodies |  |  |
| APC Mouse anti-Mouse CD45.1 (clone A20) | BD Bioscience | Cat#558701; RRID:<br>AB_1645214 |
| PerCP-Cy<sup>TM</sup>5.5 Mouse anti-Mouse CD45.2<br>(clone 104) | BD Bioscience | Cat#552950; RRID:<br>AB_394528 |
| APC-Cy<sup>TM</sup>7 Rat anti-Mouse CD45 (clone 30-F11) | BD Bioscience | Cat#557659; RRID:<br>AB_396774 |
| PerCP-Cy<sup>TM</sup>5.5 Rat anti-CD11b (clone M1/70) | BD Bioscience | Cat#550993; RRID:<br>AB_394002 |
| PE-Cy<sup>TM</sup>7 Hamster anti-Mouse CD11c (clone<br>HL3) | BD Bioscience | Cat#558079; RRID:<br>AB_647251 |
| BV 605<sup>TM</sup> anti-mouse CD11c Antibody (clone<br>N418) | BioLegend | Cat#117334; RRID:<br>AB_117334 |
| PE-Cy<sup>TM</sup>7 Rat Anti-Mouse Ly6C (clone AL-21) | BD Bioscience | Cat#560593; RRID:<br>AB_1727557 |
| FITC Rat anti-Mouse Ly6G (clone 1A8) | BD Bioscience | Cat#551460; RRID:<br>AB_394207 |
| FITC Rat anti-Mouse CD45R/B220 (clone<br>RA3-6B2) | BD Bioscience | Cat#553087; RRID:<br>AB_394617 |
| PE anti-mouse CD183 Antibody (clone<br>CXCR3-173) | BioLegend | Cat#126505; RRID:<br>AB_1027656 |
| BV421 Rat Anti-Mouse SiglecF (clone E50-2440) | BD Bioscience | Cat#562681; RRID:<br>AB_2722581 |
| BV605 Rat Anti-Mouse SiglecH (clone 440c) | BD Bioscience | Cat#747673; RRID:<br>AB_2744234 |
| Purified Rat Anti-Mouse CD16/CD32 (clone 2.4G2) | BD Bioscience | Cat#553142; RRID:<br>AB_394657 |
| eFluor 450 NK1.1 Monoclonal Antibody (clone PK136) | eBioscience | Cat#48-5941-80; RRID:<br>AB_2043878 |
| eFluor 450 CD19 Monoclonal Antibody (clone eBio1D3) | eBioscience | Cat#48-0193-82; RRID:<br>AB_2734905 |
| Alexa Fluor 700, I-A/I-E Monoclonal Antibody (clone M5/114.15.2) | eBioscience | Cat#56-53-21; RRID:<br>AB_494009 |
| FITC Mouse IgG1 kappa Isotype Control | eBioscience | Cat#11-4714-42; RRID:<br>AB_10596964 |
| PE Rat IgG2a kappa Isotype Control | eBioscience | Cat#12-4321-42; RRID:<br>AB_1518773 |
| PerCP-Cyanine5.5 IgG Isotype Control | eBioscience | Cat#45-4888-80; RRID:<br>AB_906260 |
| APC Rat IgG2b kappa Isotype Control | eBioscience | Cat#17-4031-82; RRID:<br>AB_470176 |
| APC-eFluor 780 Rat IgG2b kappa Isotype Control | eBioscience | Cat#47-4031-80; RRID:<br>AB_1272021 |
| Alexa Fluor 700 Rat IgG2b kappa Isotype Control | eBioscience | Cat#56-4031-80; RRID:<br>AB_837123 |
| PE-Cyanine7 Rat IgG2a kappa Isotype Control | eBioscience | Cat#25-4321-82; RRID:<br>AB_470200 |
| eFluor 450 Rat IgG1 kappa Isotype Control | eBioscience | Cat#48-4301-82; RRID:<br>AB_1271984 |
| Bacterial and Virus Strains |  |  |
| AF293 | Fungal genetics stock center | #A1100 |
| AF293 ds-Red | (20) | N/A |
| CEA10 (also known as CBS144.89) | (47) | Received from Robert A. Cramer |
| Chemicals, Peptides, and Recombinant Proteins |  |  |
| Alexa Fluor 633 succinimidyl ester | Invitrogen | Cat#S12375 |
| Collagenase IV | ThermoFisher Scientific | Cat#LS004189 |
| RPMI-1640 | RPMI-1640 | 21870092 |
| Enrofloxacin | Bayer | Baytril 100 |
| Voriconazole | Pfizer | N/A |
| Tween-20 | Sigma | P9416 |
| Protease Inhibitor Cocktail | Roche | Cat#11697498001 |
| DNase I, Grade II | Sigma | Cat#10104159001 |
| isoflurane | Henry Schein Animal Health | Cat#29405 |
| Diphtheria toxin (DT) | List Biological Laboratories | Cat#150 |
| Paraformaldehyde, 32% Solution | ThermoFisher Scientific | Cat#50-980-495 |
| CM-H<sub>2</sub>DCFDA | ThermoFisher Scientific | Cat#C6827 |
| 10× HBSS | ThermoFisher Scientific | 14065056 |
| TRIzol-LS | Invitrogen | 10296028 |
| Mouse IFNα2/4 ELISA kit | Invitrogen | BMS6027 |
| Mouse IL-28B/IFN-lambda 3 DuoSet ELISA | Invitrogen | DY1789B-05 |
| Mouse GM-CSF DuoSet ELISA | Invitrogen | DY415-05 |
| Mouse TNF alpha ELISA Ready-SET-Go!<sup>TM</sup> Kit | Invitrogen | 50-173-31 |
| Mouse IL-6 ELISA Ready-SET-Go!<sup>TM</sup> Kit | Invitrogen | 50-172-19 |
| Mouse IL-1 beta ELISA Kit | Invitrogen | BMS6002 |
| Mouse IL-12 p70 ELISA Kit | Invitrogen | BMS6004 |
| High-Capacity RNA-to-cDNA Kit | Appliedbiosystems | 4387406 |
| Ribosomal RNA Control Reagents | Appliedbiosystems | 4308329 |
| IFNa2 TaqMan Assay Mm00833961_s1 | Thermo Fisher | 4331182 |
| IFNa1/a5/a6 TaqMan Assay<br>Mm03030145_gH | Thermo Fisher | 4331182 |
| IFNL2/3 TaqMan Assay Mm04204155_gH | Thermo Fisher | 4331182 |
| Experimental Models: Organisms/Strains |  |  |
| BDCA2-DTR | The Jackson Laboratory | JAX: 014176 |
| C57BL/6 | The Jackson Laboratory | JAX: 000664 |
| C57BL/6.SJL | Taconic | Stock No. 4007 |
| *Stat1*<sup>-/-</sup> mice | The Jackson Laboratory | JAX: 012606 |
| *GBP*<sup>chr3-/-</sup> mice |  | Received from Dr. Thirumala-Devi Kanneganti. |
| *Cybb*<sup>-/-</sup> mice | The Jackson Laboratory | JAX: 002365 |
| Software and Algorithms |  |  |
| Prism 10 | Prism 10 | N/A |
| Flow Jo 10.10.0 | Flow Jo 10.10.0 | N/A |

