## Supplementary figures and images for "An IFN-STAT1-CYBB Axis Defines Protective Plasmacytoid DC to Neutrophil Crosstalk During *Aspergillus fumigatus* Infection"

### Related to Fig. 1

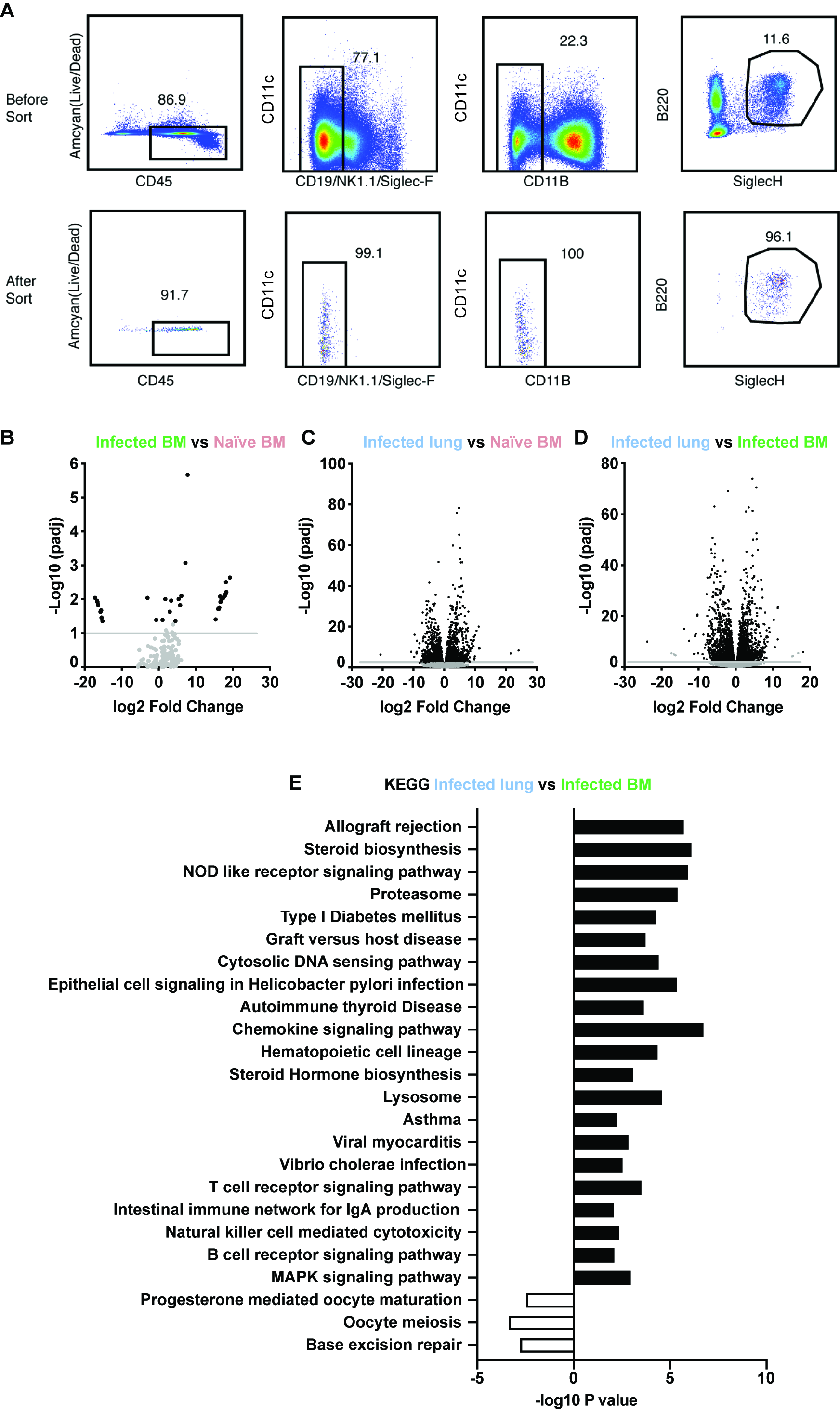

### Related to Fig. 3

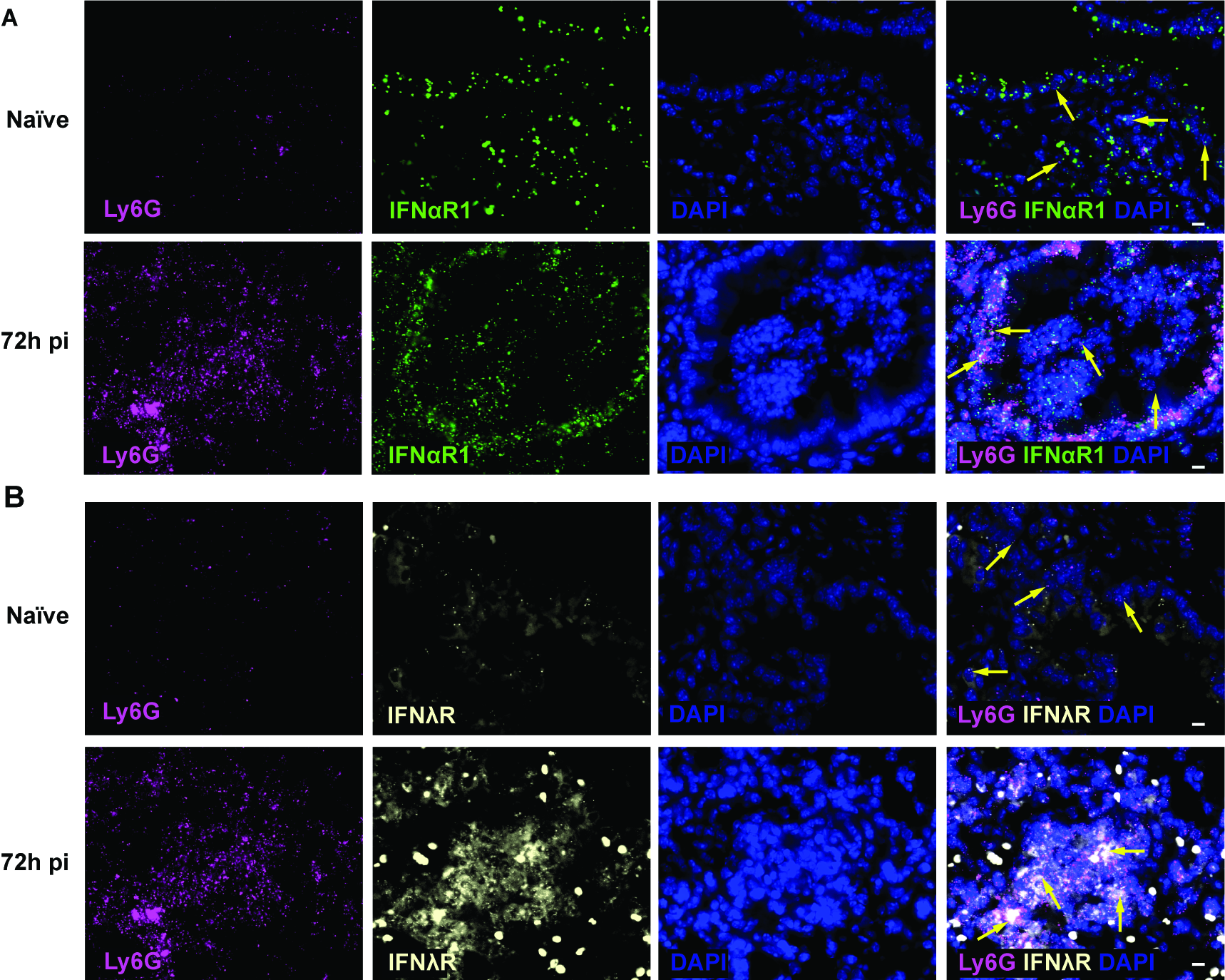

### Related to Fig. 4

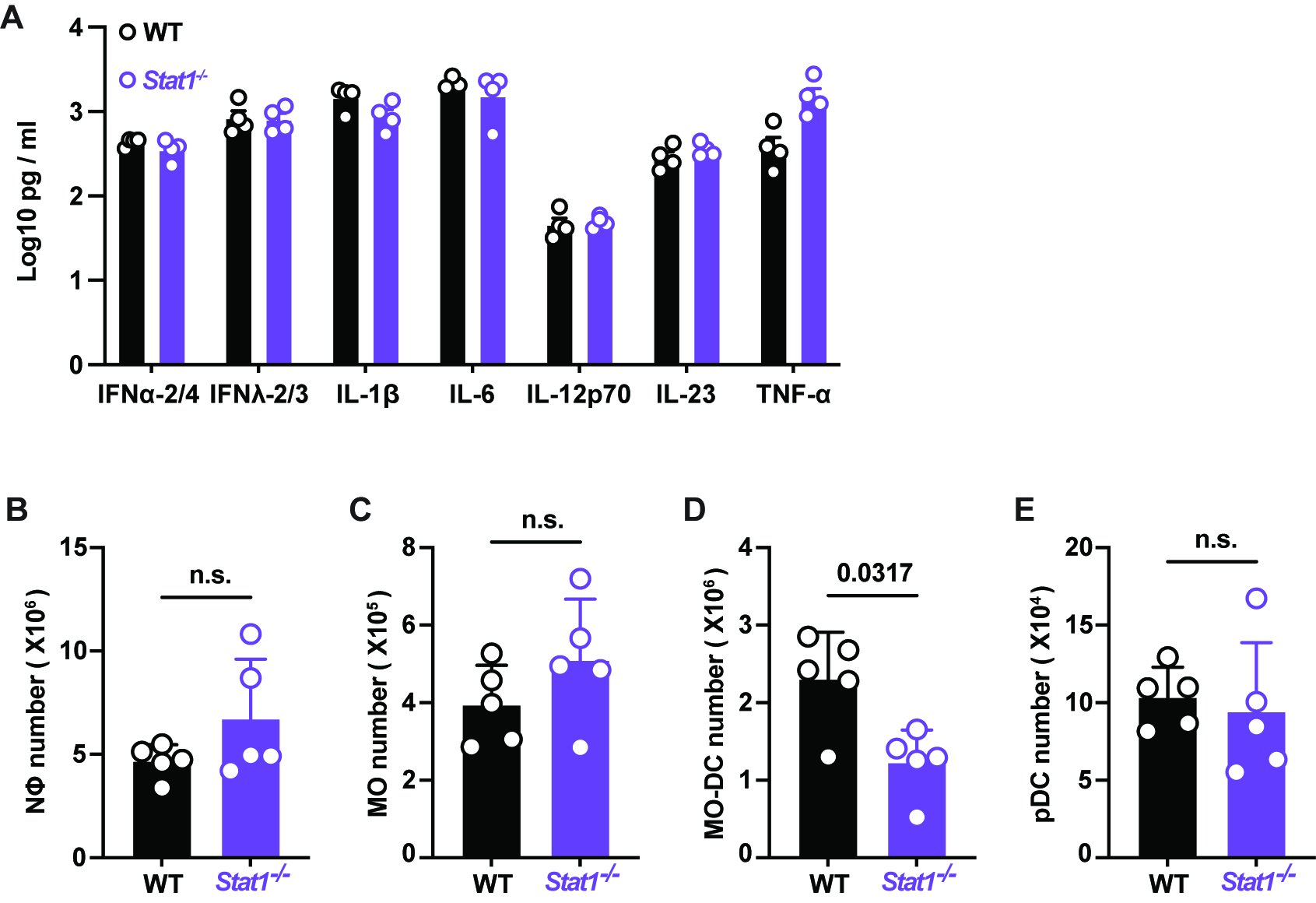

### Related to Fig. 6

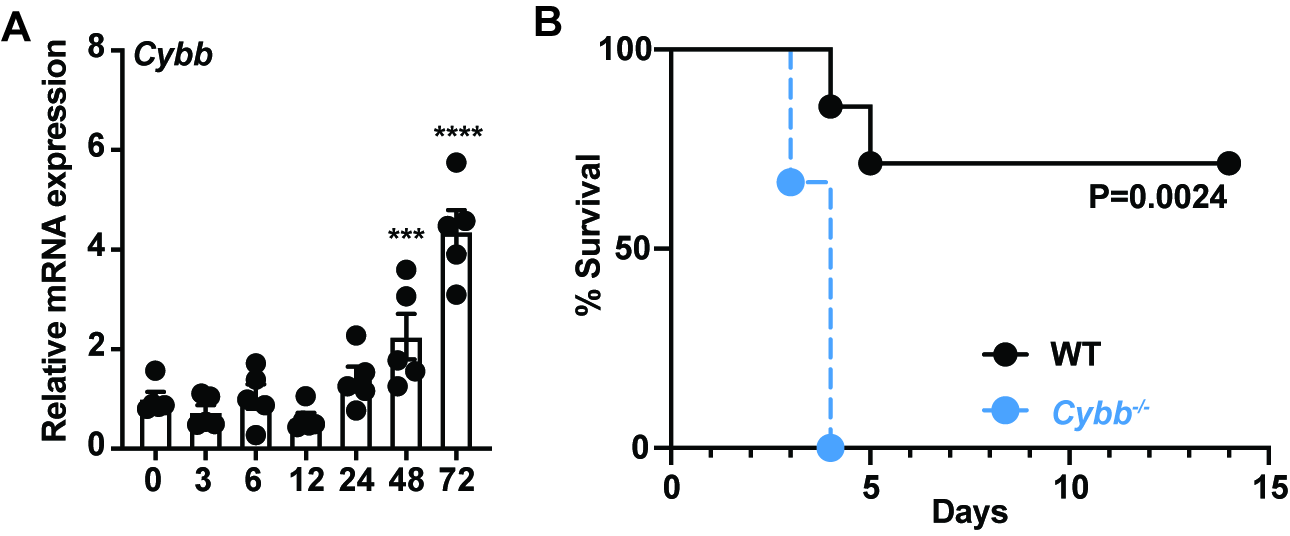
